# Diffusion Neuroimaging of Speech Acquisition in Infants

**DOI:** 10.1101/2025.05.23.655876

**Authors:** Feihong Liu, Jiameng Liu, Jinchen Gu, Yaoxuan Wang, Xinyi Cai, Zhaoyan Wang, Hao Wu, Jun Feng, Han Zhang, Dinggang Shen

**Affiliations:** School of Biomedical Engineering & State Key Laboratory of Advanced Medical Materials and Devices, ShanghaiTech University, Shanghai, 201210, China; School of Information and Technology, Northwest University, Xi’an, 710127, China; Department of Otolaryngology-Head and Neck Surgery, Shanghai Ninth People’s Hospital, Shanghai Jiao Tong University School of Medicine, Shanghai, 200011, China; Ear Institute, Shanghai Jiao Tong University School of Medicine, Shanghai, 200011, China; Shanghai Key Laboratory of Translational Medicine on Ear and Nose Diseases, Shanghai, 200011, China; Shanghai Clinical Research and Trial Center, Shanghai, 201210, China; Shanghai United Imaging Intelligence Co., Ltd., Shanghai, 200230, China

## Abstract

How multisystem white-matter maturation supports early speech acquisition in infancy remains unclear. Using longitudinal diffusion MRI from the Baby Connectome Project, we quantified neurite density maturation from birth to 24 months across 33 white-matter tracts spanning motor, auditory, visual, ventral-language, and dorsal-language systems. Maturation was coordinated across systems rather than confined to canonical language pathways: dorsal-language association tracts began relatively immature near birth but showed steep postnatal growth, progressively rebalancing with motor and sensory systems. Tract–behavior coupling was likewise distributed across developmental domains, with broadly positive age-adjusted associations for motor outcomes, a sparser, partly motor-like profile for gesture-related phrase comprehension, and predominantly negative associations for lexical speech-related outcomes. Exploratory follow-up analyses further suggested motor and social precursor capacities that may help mediate these indirect developmental routes from structural maturation to later speech-related outcomes. These findings support a multisystem developmental account in which early speech acquisition emerges not from an isolated language pathway, but from coordinated maturation across multiple systems and the precursor skills that link them.

## 1 Introduction

### A multisystem developmental framework for speech acquisition

Spoken communication emerges during infancy, amid rapid neural maturation and concurrent gains in motor control, perception, and social engagement (*1,2*). From birth to the first year of life, infants progress from cries and other arousal-related behaviors to increasingly structured vocal, gestural, and word-like forms of communication (*3, 4, 5*), while simultaneously refining speech perception and engaging in reciprocal exchanges with caregivers (*6, 7, 8*). In this developmental sense, we use *early communication* to refer to the broader developmental pathway within which speech acquisition unfolds, including gesture- and action-based precursor behaviors that are theoretically and empirically linked to later spoken language (*8*). Yet it remains unclear how neural maturation supports this developmental cascade, and whether such support is localized to canonical language pathways or distributed across multiple brain systems.

This developmental transition is unlikely to depend on a single language-specific pathway (*9,10*). Instead, increasing evidence suggests that speech acquisition involves a multisystem process in which auditory input is coordinated with motor control, visual cues, and social engagement (*11, 12, 13*). Such coordination should be especially important in infancy, when mappings between variable sensory input and stable communicative categories are still being established (*14, 15*). The central question is therefore not merely whether a particular language pathway predicts later language outcomes, but how multiple brain systems mature in concert to support the progression from simple precursor behaviors to spoken communication.

White matter provides a natural target for this question because it links distributed brain regions and supports information exchange across functionally segregated networks, thereby offering a system-level view of developmental brain reorganization (*16, 10*). Although diffusion MRI (dMRI) provides a non-invasive window onto white-matter microstructural maturation across infancy, prior infant MRI studies have typically emphasized a small number of dorsal or ventral language tracts, relatively modest samples, or focused speech-behavior assessment (*17, 11, 18*). Consequently, it remains unclear how language-relevant pathways co-develop with auditory, visual, and motor tracts, or whether tract–behavior coupling is concentrated within canonical language domains or distributed across multiple developmental domains. Addressing these questions is technically challenging, when rapid tissue change and age-related contrast shifts during the first 24 months of life complicate infant neuroimaging data processing and anatomically plausible tract extraction (*19, 20, 21, 22*).

### Sensory–motor coordination as a candidate developmental mechanism

Within this multisystem framework, sensory–motor coordination provides a plausible developmental mechanism through which early communication may unfold as a cascading process during infancy (*23, 13*). Here, *sensory* refers primarily to the auditory and visual pathways through which speech-related input is encoded, whereas *motor* refers to the action systems that support communicative behavior, including vocal, orofacial, gestural, and other action-based responses (*24, 25*). From a sensorimotor perspective, infants do not acquire speech by passively encoding sounds alone (*24*). Rather, they progressively learn correspondences among heard speech, seen articulatory cues, and their own communicative actions, with these mappings further stabilized through social interaction (*26, 27, 28, 29, 30*). Auditory, visual, and motor systems therefore constitute a coordinated substrate through which early arousal-related behaviors can develop into increasingly structured forms of communication (*24, 13*).

Evidence for such coordination appears early, even before birth. Fetuses show motor and autonomic responses to acoustic stimulation, including fetal movements and changes in heart rate (*31, 32, 33, 34*), and fetal dMRI studies have identified long-range association tracts, including the arcuate fasciculus (AF) and superior longitudinal fasciculus (SLF), linking temporal auditory regions with inferior frontal cortex (IFC; approximately BA44 within the inferior frontal gyrus [IFG]) (*35, 36*). After birth, infants detect correspondences between heard speech and seen articulations, including audiovisual matching of speech sounds and facial movements, as well as sensitivity to audiovisual synchrony (*9, 37, 38*). Visual and motor speech cues can further facilitate phoneme discrimination and speech perception when auditory input is noisy or otherwise ambiguous (*39, 40, 41*); and infants show increased activation in motor cortical regions when perceiving unfamiliar non-native speech relative to familiar native speech stimuli (*27*). Social experience adds other components: live, contingent interpersonal interaction supports phonetic learning more effectively than matched video- or audio-only exposure, caregiver feedback guides the development of prelinguistic vocalizations, and combined sensory–motor experience appears to support learning more effectively than sensory or motor experience alone (*42, 43, 44, 30*).

Taken together, current evidence suggests that early communication develops by progressively linking sensation, action, and social contingency, rather than through the isolated maturation of a single tract or through auditory experience alone. This framework yields a tract-level prediction: if speech acquisition is embedded in a sensory–motor developmental cascade, then coordinated maturation should be detectable across auditory, visual, motor, and language-related white-matter systems, and tract–behavior coupling would likewise extend across multiple developmental domains.

To test this account, we designed a hierarchical analysis framework (Figure 1) leveraging the Baby Connectome Project (BCP), a large longitudinal infant neuroimaging and behavioral cohort (*45*), together with infant-oriented tract extraction methods (*21, 46, 22*). Using neurite orientation dispersion and density imaging (NODDI) (*47*), we quantified neurite density maturation from birth to 24 months across 33 white-matter tracts spanning five systems — auditory, visual, motor, ventral-language, and dorsal-language (Figure 2). We then related tract microstructure to 11 developmental assessments spanning six domains — speech-related outcomes, motor, social interaction, visual/cognitive, daily living, and emotion/regulation (Table 1). Within this framework, tract maturation trajectories and tract–behavior associations constitute the core analyses of the manuscript, whereas cross-sectional prioritization of candidate developmental intermediates and time-ordered longitudinal mediation models are treated as targeted exploratory follow-up analyses. This design allowed us to test whether early communication is better understood as an emergent property of coordinated maturation across multiple functionally segregated yet developmentally coordinated brain systems than as a process attributable to any single canonical language pathway or developmental domain.

**Figure 1:**
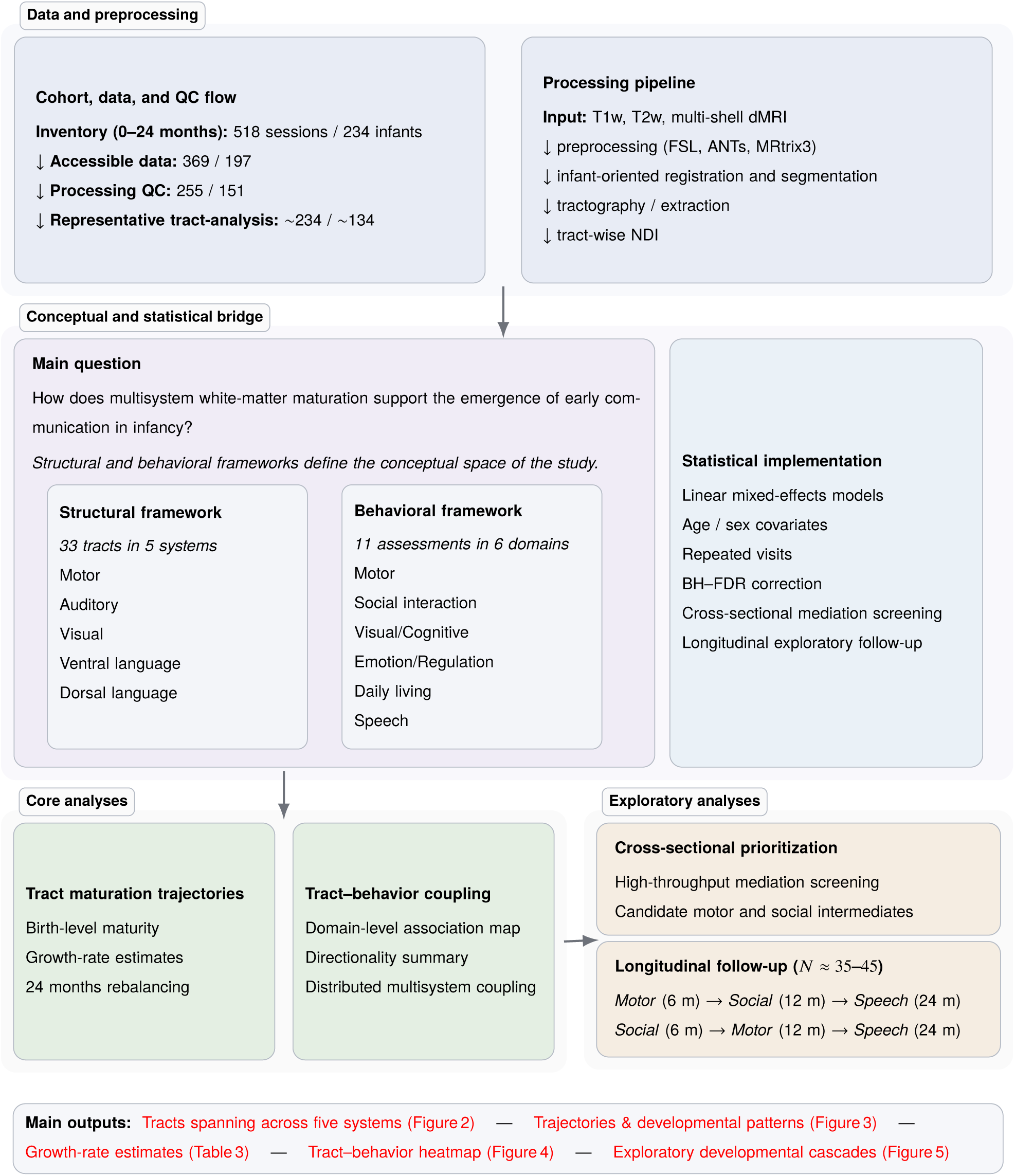
Study overview and analytic framework. Leveraging the BCP cohort, we retained processed diffusion MRI data that passed the quality-control pipeline and organized subsequent analyses within a hierarchical framework. The framework included 33 tracts spanning five systems, whereas the behavioral framework comprised 11 assessments spanning six developmental domains (Table 1). Core analyses focused on tract maturation trajectories and tract–behavior coupling, followed by exploratory cross-sectional prioritization and longitudinal follow-up analyses. The bottom band summarizes the principal outputs reported in subsequent figures and tables.

**Figure 2:**
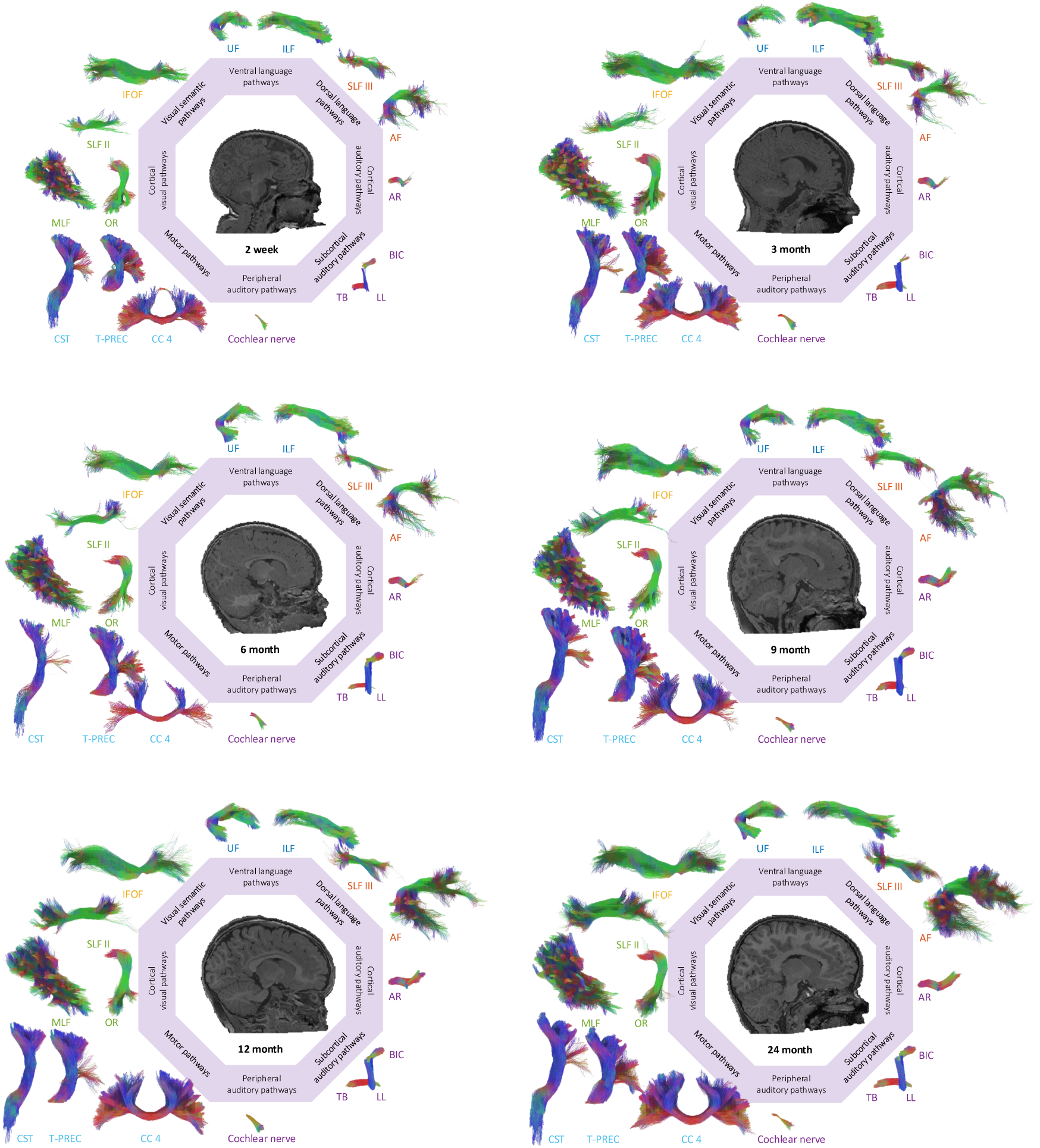
White-matter tract framework for the representative left-hemisphere analyses. Tract renderings are shown from the neonatal period to 24 months and arranged using an octahedral layout schematic to compactly display multiple developmental stages within a single figure. This layout is used for visual organization only and does not encode diffusion sampling directions, HARDI orientation structure, or tract directionality. Tracts are grouped by text color into five systems: motor (CST, T-PREC, CC4), auditory (Cochlear nerve, TB, LL, BIC, AR), visual (OR, MLF, SLF II), ventral-language (UF, ILF, IFOF), and dorsal-language (AF, SLF III). IFOF is highlighted separately in yellow because of its dual relevance to ventral-language and visual–semantic processing. This representative tract set is used for developmental trajectory and maturity-structure summaries in Figure 3; tract–behavior analyses use the full 33-tract framework in Figure 4. CST: corticospinal tract; T-PREC: thalamo-precentral; and CC4: corpus callosum anterior midbody (primary motor); TB: trapezoid body; LL: lateral lemniscus; BIC: brachium of the inferior colliculus; AR: acoustic radiation; OR: optic radiation; SLF II: superior longitudinal fasciculus II; MLF: middle longitudinal fasciculus; UF: uncinate fasciculus; ILF: inferior longitudinal fasciculus; IFOF: inferior fronto-occipital fasciculus; AF: arcuate fasciculus; SLF III: superior longitudinal fasciculus III.

**Table 1:**
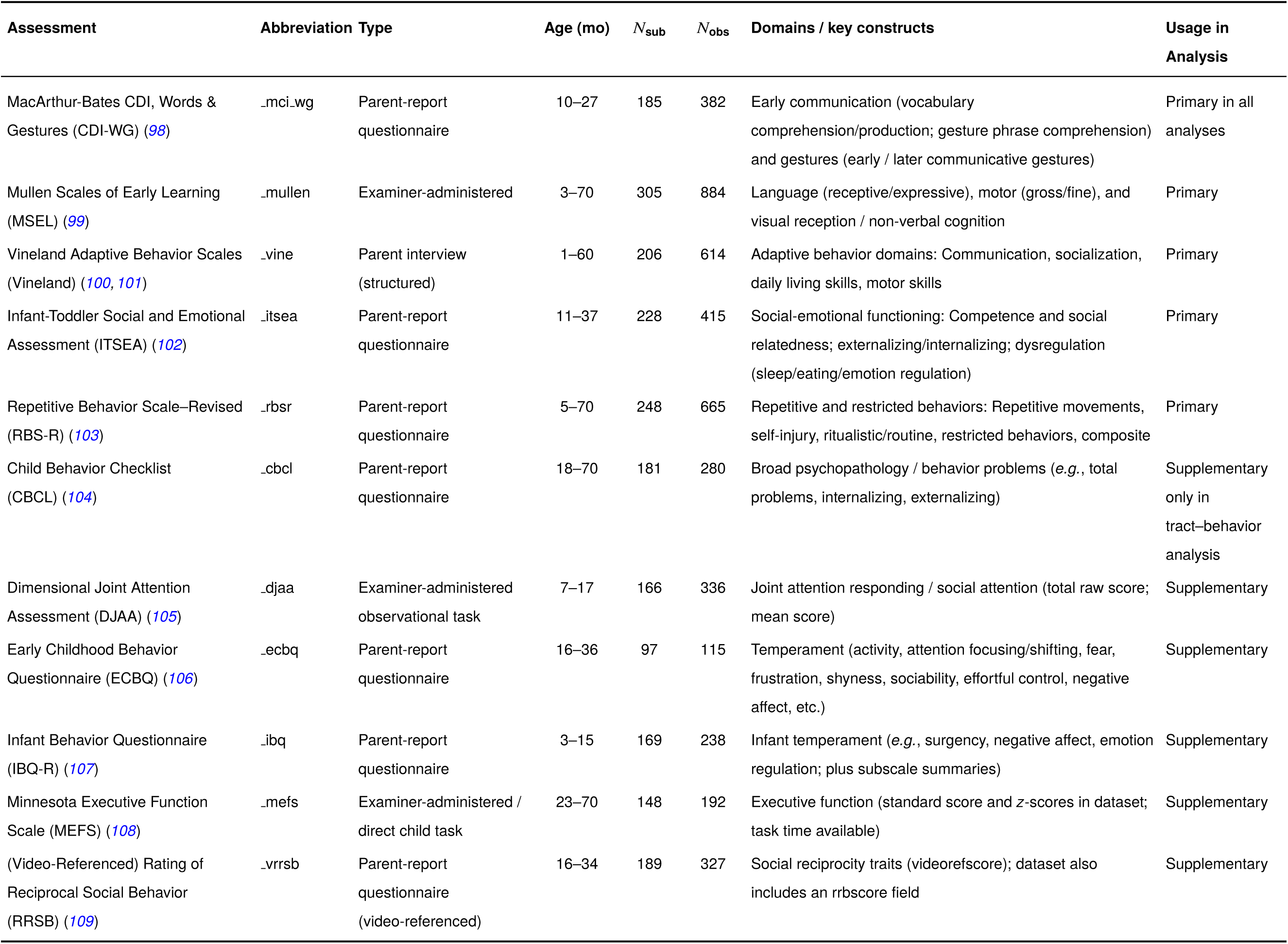
Summary of eleven behavioral assessments used in the study. Age indicates the observed interview age in months (min–max) in the BCP cohort. *N*_sub_ indicates the number of participants with non-missing interview age for that instrument with *N*_obs_ indicating the number of visits in total. “*Primary*” denotes instruments included in all analyses, whereas “*Supplementary*” denotes measures collected for a subset of participants which are only used in the tract–behavior analyses. By combining multiple instruments, we assessed six developmental domains (Motor skills, social interaction, visual/cognitive Skills, daily living skills, emotional/regulatory behavior, and communication/speech), as referenced in the manuscript.

## 2 Results

### Cohort, data processing, and tract system

We first established the cohort, diffusion MRI processing pipeline, tract system, and behavioral framework used in all downstream analyses. Figure 1 summarizes the overall study design, from cohort derivation and data processing to the distinction between core analyses and exploratory follow-up analyses. Within the 0–24-month BCP imaging inventory, 518 scan sessions from 234 infants were identified. Of these, 369 scan sessions from 197 infants had accessible imaging data, and 255 scan sessions from 151 infants were retained after processing quality control. A representative tract-analysis sample comprised approximately 234 scan sessions from approximately 134 infants, with exact denominators varying modestly across tract- and behavior-specific models (Table 2).

**Table 2:**
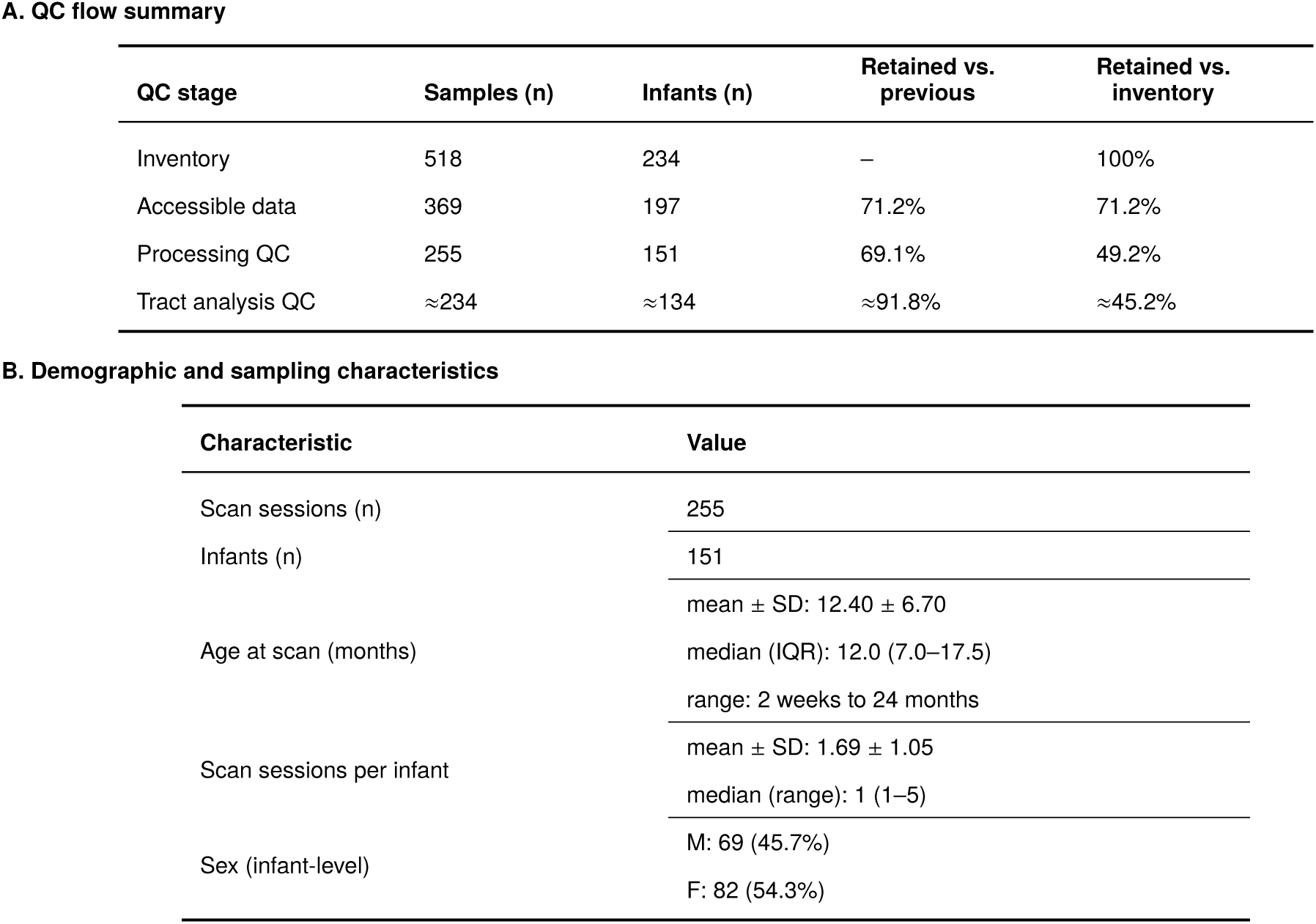
Cohort derivation and characteristics of the processing-QC sample. Panel A summarizes the quality-control flow across analytic stages. Panel B summarizes demographic and sampling characteristics of the processing-QC cohort. Tract-analysis counts are representative rather than universal, because exact denominators vary modestly across tract- and behavior-specific models.

The processing-QC cohort spanned 2 weeks to 24 months of age (mean 12.40 ± 6.70 months; median 12.0, IQR 7.0–17.5), and infants had a mean of 1.69 ± 1.05 scan sessions (median 1, range 1–5; Table 2). Diffusion MRI data were processed through an infant-oriented tract pipeline to derive tract-wise neurite density index (NDI), the primary microstructural metric used throughout the study. Across downstream analyses, we examined 33 tracts spanning five systems — auditory, visual, motor, ventral-language, and dorsal-language — and related them to 11 assessments spanning six developmental domains — speech-related outcomes, motor, social interaction, visual/cognitive, daily living, and emotion/regulation (Figure 2; Table 1). For developmental trajectory and maturity-structure summaries, we present a representative left-hemisphere tract set spanning these systems, whereas tract–behavior analyses use the full 33-tract framework. For visualization of tract–behavior coupling, speech-related outcomes were further displayed as lexical and gesture measures to distinguish broader lexical outcomes from earlier, more action-linked communicative behavior.

### Multisystem tract maturation and progressive cross-system reorganization

The representative tract system is shown in Figure 2, spanning auditory, visual, motor, ventral-language, and dorsal-language systems from the neonatal period to 24 months. Within this system, tracts differed substantially in their developmental profiles. Ventral pathways such as ILF and UF showed comparatively slower maturation, whereas several dorsal language tracts matured more rapidly across the first two postnatal years. In the neonatal stage, the left AF terminated predominantly in precentral cortex, with sparse but detectable extension toward inferior frontal cortex (BA44; Figure S4), consistent with predominant premotor connectivity at birth (*48, 49*) while suggesting a weak early structural scaffold toward inferior frontal territory.

We next quantitatively characterized tract maturation across the representative left-hemisphere tracts (Figure 3). Model-predicted NDI trajectories increased across infancy for all tracts (Eqn. 2), but they differed markedly in both their near-birth maturity structure and subsequent developmental profiles (Figure 3 **a**). At 2 weeks, maturity was highly asynchronous across systems (Figure 3 **b**): several auditory and motor tracts occupied the upper end of the predicted NDI range, whereas AF, SLF II, and SLF III were among the least mature tracts.

**Figure 3:**
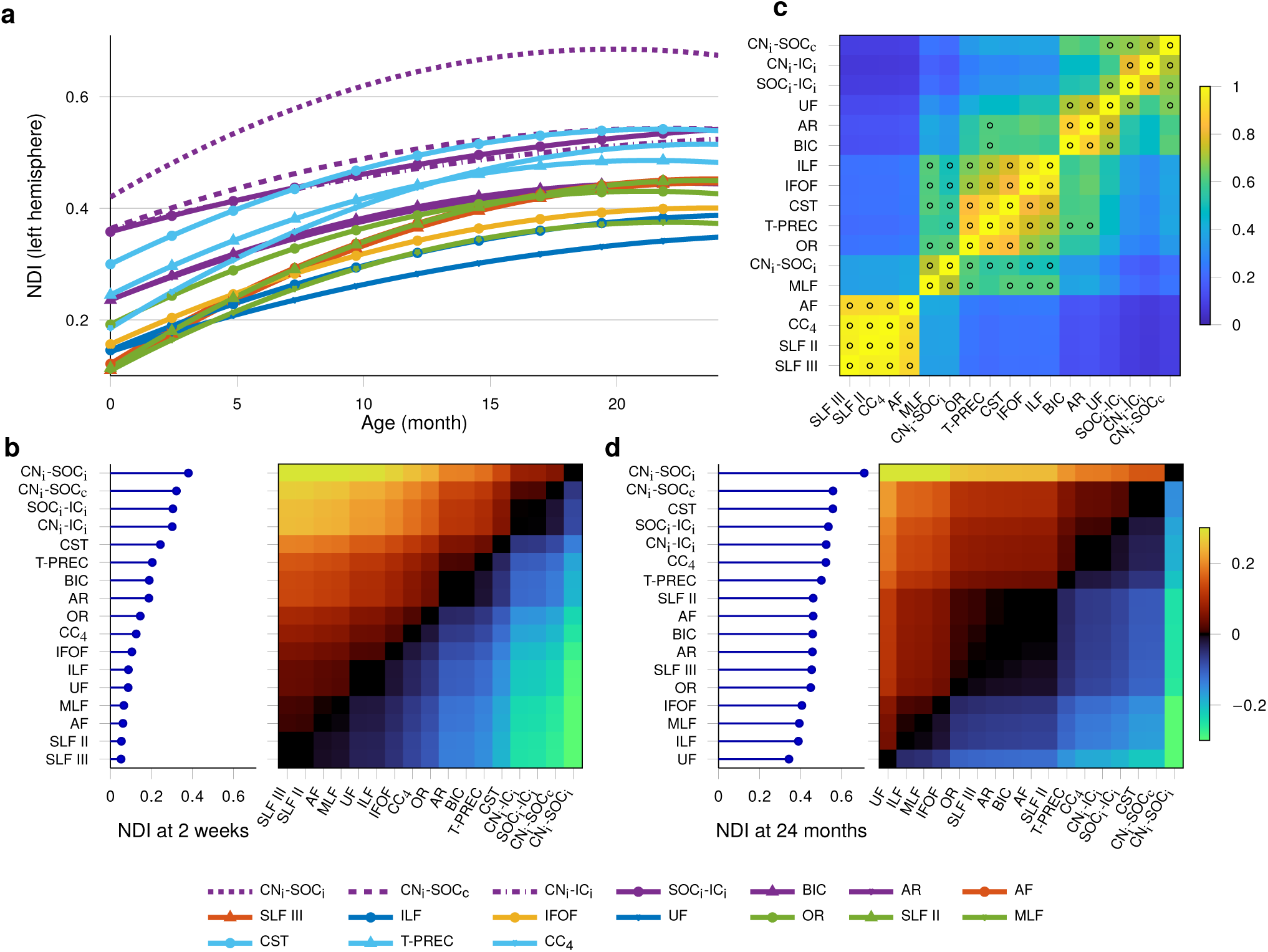
Tract maturation trajectories and model-predicted maturity pattern across the 17 representative left-hemisphere tracts. (**a**) Model-predicted developmental trajectories from birth to 24 months. (**b**) Near-birth maturity pattern, shown as tract rank and a signed pairwise difference matrix at 2 weeks, where each cell equals the predicted NDI of the row tract minus that of the column tract; values near 0 indicate more similar maturity levels. (**c**) Trajectory-shape similarity, with overlaid markers denoting tract pairs whose logAge24 slopes did not significantly differ after BH–FDR correction across all 136 pairwise slope comparisons within the displayed representative tract set, based on the joint mixed-effects model fit across tract. (**d**) Twenty-four-month maturity pattern, shown as tract rank and a signed pairwise difference matrix, defined analogously to panel **b**. Panels **b** and **d** share the same signed-difference color scale; the colorbar shown beside **d** applies to both pairwise-difference matrices. i: ipsilateral; c: contralateral.

Despite these immature starting levels, several association tracts showed rapid subsequent growth. AF, SLF II, SLF III, and CC_4_ displayed closely aligned developmental trajectories over the first 24 months (Figure 3 **a**), and trajectory-shape similarity analysis (Eqn. 6) highlighted a high-similarity cluster among these tracts (Figure 3 **c**). For visualization, overlaid markers further identified tract pairs for which the pairwise logAge_24_ slopes did not significantly differ after BH–FDR correction. These markers are used as descriptive evidence of statistically indistinguishable maturation-rate estimates, from the joint mixed-effects model fit across 136 pairwise slope comparisons within the displayed 17 representative tract set (Eqn. 7). It supported the interpretation that AF, SLF II, SLF III, and CC_4_ form a coherent maturation group rather than merely appearing visually similar, and other high-similarity clusters also indicated coordinated early development across systems.

Single-tract log-age mixed-effects models (Eqn. 3) further quantified their rapid postnatal growth (Table 3). The largest absolute maturation-rate estimates were observed for SLF II_*L*_ (*β*_log *Age*_ = 0.1531), SLF III_L_ (0.1520), CC_4_ (0.1496), and AF_L_ (0.1493), whereas lower estimates were observed for several ventral, sensory, and auditory tracts, including UF_L_ (0.0934), AR_L_ (0.0951), BIC_L_ (0.0952), SOC_L_–IC_L_ (0.0827), and CN_L_–IC_L_ (0.0783). Thus, AF, SLF II, and SLF III were characterized not by high near-birth maturity, but by relatively steep postnatal growth.

**Table 3:** Tract-wise maturation-rate estimates from single-tract log-age mixed-effects models. Entries report tract-specific fixed-effect slopes for log(1 + *Age*) from models fit separately to each representative left-hemisphere tract (Eqn 3). Larger values indicate steeper age-related increases in NDI and are used here as compact summaries of relative postnatal maturation rate.

|  |  |  |  |  |  |  |  |  |  |
| --- | --- | --- | --- | --- | --- | --- | --- | --- | --- |
| SLF II | SLF III | CC<sub>4</sub> | AF | MLF | CST | ILF | IFOF | OR | T-PREC |
| 0.1531 | 0.1520 | 0.1496 | 0.1493 | 0.1196 | 0.1090 | 0.1075 | 0.1060 | 0.1051 | 0.1008 |
| CN<sub>i</sub>–SOC<sub>i</sub> | BIC | AR | UF | CN<sub>i</sub>–SOC<sub>c</sub> | SOC<sub>i</sub>–IC<sub>i</sub> | CN<sub>i</sub>–IC<sub>i</sub> |  |  |  |
| 0.1216 | 0.0952 | 0.0951 | 0.0934 | 0.0830 | 0.0827 | 0.0783 |  |  |  |

*Note.* These values correspond to the tract-wise absolute maturation-rate estimates from the single-tract models based on $\log(1 + \text{Age})$ .

By 24 months, the tract maturity pattern had become more balanced (Figure 3 d). Pairwise signed differences showed that, though projection tracts such as CST remained distinct, AF, SLF II, and SLF III approached the maturity range of primary sensory tracts such as AR and OR. We therefore interpreted the structural pattern as progressive cross-system rebalancing rather than as a simple contrast between initial immaturity and rapid later growth. Complementary supplementary analyses were consistent with this interpretation at the system level: inter-tract dispersion (coefficient of variation, as defined in Eqn. S1) decreased with age, and motor-leading sensory–motor gradients showed more associations with association tracts and with motor behavioral outcomes than did sensory–sensory contrasts (Figure S13,Table S6, and S7).

### Distributed tract-behavior coupling across developmental domains

We next tested whether tract microstructure covaried with developmental skills across the full 33 tract system. After Benjamini–Hochberg false discovery rate (BH–FDR) correction across the 33 tracts within each behavioral variable (*50*), significant tract–behavior associations were widely distributed across all five tract systems and six developmental domains rather than confined to canonical language tracts or auditory domain alone (Figure 4). Complete lists of retained associations are provided in the Supplementary Data 2 and Supplementary Data 3. Because speech-related measures were split for visualization into lexical and gesture displays, the heatmap shows seven columns derived from the underlying six-domain framework (Figure 4).

**Figure 4:**
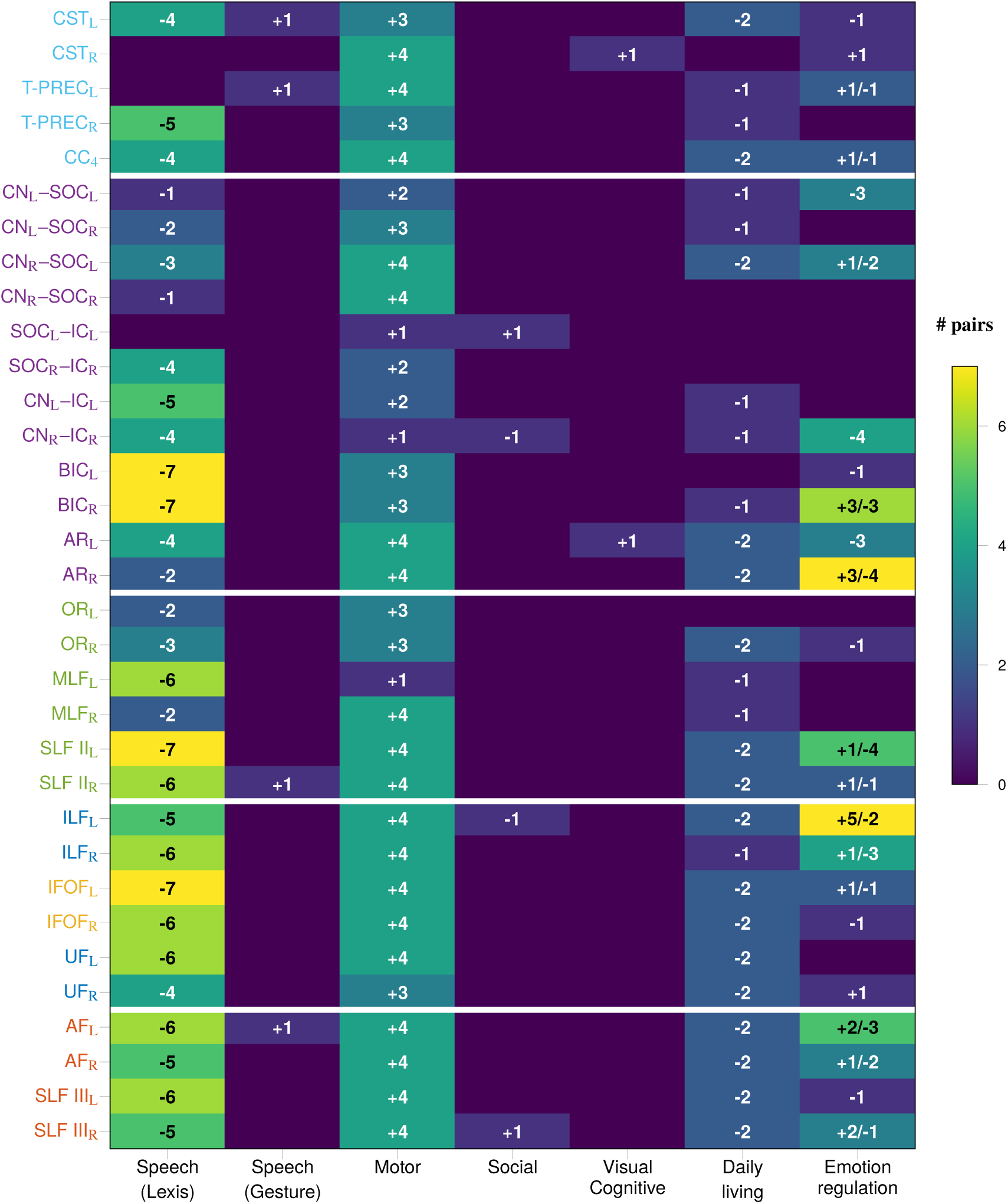
Domain-level tract–behavior associations across the full 33-tract framework. Rows list white-matter tracts grouped by system (white separators; text colors indicate motor, auditory, visual, ventral-language, and dorsal-language systems). Columns summarize six developmental domains. For visualization, speech-related outcomes are split into lexical and gesture-related subparts, yielding seven columns from an underlying six-domain behavioral framework. Cell color indicates the number of significant tract–behavior associations within each tract-by-domain cell after BH–FDR correction across the 33 tracts within each behavioral variable (*q* < 0.05). Overlaid text summarizes the directionality of the linear age-adjusted NDI component: for example, “−4” indicates four negative associations, whereas “+*a*/−*b*” indicates mixed directions across outcomes within that cell. Because the mixed-effects models included both linear and quadratic NDI terms, the sign shown here should not be interpreted as a purely monotonic developmental effect. At the domain level, motor outcomes showed predominantly positive associations, whereas lexical speech-related outcomes showed widespread negative associations indicating timing-sensitive age-adjusted coupling rather than evidence that greater white-matter maturation impairs communication.

Three patterns were especially prominent. First, motor outcomes showed predominantly positive associations distributed across auditory, visual, motor, ventral-language, and dorsal-language tracts, indicating that greater tract maturity broadly covaried with stronger age-adjusted motor performance. Second, lexical speech-related outcomes showed widespread predominantly negative associations across the same tract system. Third, gesture-related measures showed a distinct, sparser, and partly motor-like profile rather than the broadly negative pattern observed for lexical speech outcomes.

Other developmental domains showed more selective or heterogeneous associations. Visual/cognitive and daily living measures tended to show internally consistent association directions, whereas social-interaction and emotion/regulation measures showed more mixed tract-level effects. The sign displayed in the heatmap reflects the direction of the linear age-adjusted NDI term, not necessarily a purely monotonic developmental effect. In particular, the predominantly negative speech-related linear coefficients should not be interpreted as evidence that greater white-matter maturation impairs communication; rather, they are more consistent with timing-sensitive multisystem coupling during a period in which *neural substrates, precursor capacities, and communication outcomes may be developing both synchronously and asynchronously*.

### Exploratory developmental cascades linking motor, social, and speech-related measures

To organize the distributed tract–behavior associations, we conducted exploratory mediation analyses focused on candidate motor and social intermediate constructs in a developmental pathway. Detailed cross-sectional results are provided in the Supplementary Data 4, and pathway families are summarized in Supplementary Table S8.

Figure 5 **a** summarizes the high-throughput cross-sectional screen. Of 74,409 specified triplets, 74,276 were testable and 849 were retained after study-wide BH–FDR correction. Among these retained triplets, 118 involved a tract predictor and 731 involved a behavioral predictor (Figure 5 **b**), indicating that behavior-to-behavior coordination itself constituted a major component of the retained developmental structure. Within the retained tract-predictor triplets, motor mediators strongly predominated (110/118, 93.2%). Restricting to the subset most directly relevant to the present neu- rodevelopmental hypothesis — *tract* → *Any* → *speech* — 52 retained triplets remained, of which 44 (84.6%) involved motor mediators. By contrast, social-interaction, emotional/regulation, or daily-living mediators were rare in retained tract-to-speech pathways.

**Figure 5:**
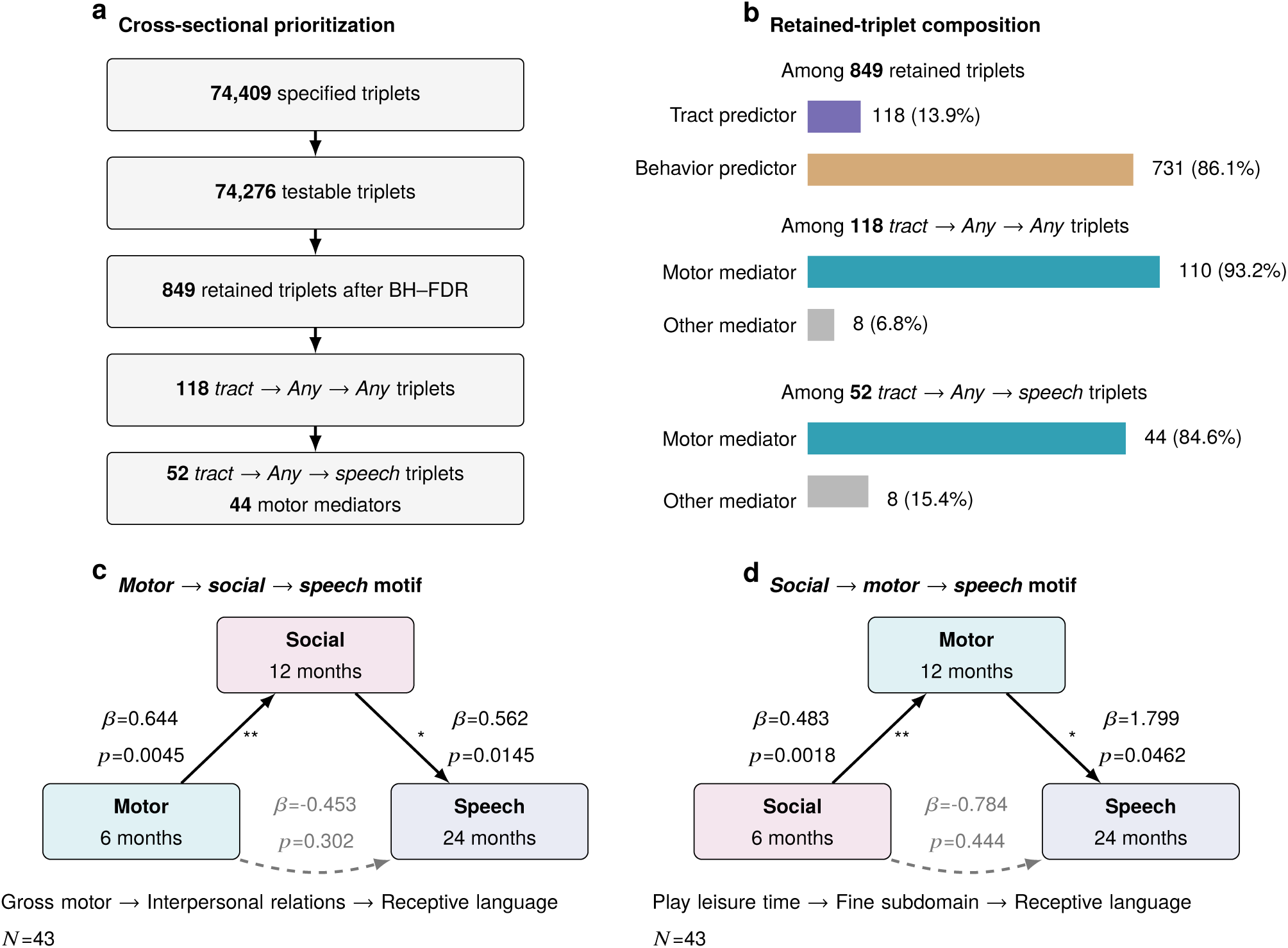
Exploratory developmental cascades linking motor, social, and speech-related measures. (**a**) Summary of the high-throughput cross-sectional mediation screen, showing the numbers of specified, testable, and retained triplets after study-wide BH–FDR correction. (**b**) Composition of retained triplets, including the proportions with tract predictors versus behavioral predictors and the predominance of motor mediators among retained tract-to-speech pathways. (**c** and **d**) Representative time-ordered longitudinal motifs in the *motor* → *social* → *speech* and *social* → *motor* → *speech* directions, respectively. Effect estimates and *p*-values shown in panels **c** and **d** are uncorrected and are provided to illustrate representative developmental motifs rather than to establish a single causal pathway. Because the longitudinal analyses were conducted in modest complete-data subsets, all longitudinal findings are interpreted as exploratory and hypothesis-generating.

When behavioral variables served as predictors, retained pathways again suggested that motor and social constructs help organize later speech-related development. Motor- or social-predictor models yielded retained pathways through multiple downstream domains, with the largest pathway counts observed for daily living, motor, and social-interaction intermediates, and fewer pathways through visual/cognitive or emotion/regulation measures (Table S8). Taken together, the cross-sectional screen suggests that tract microstructure most often links to later speech-related outcomes through motor intermediates, while broader behavior-to-behavior coordination is also substantial.

We next conducted targeted time-ordered follow-up analyses in modest complete-data subsets (*N* ≈ 35–45). Representative examples of both candidate cascades are shown in Figure 5 **c**–**d**. In the *motor* → *social* → *speech* example, early (4–6 months) gross motor ability related positively to later (12–14 months) interpersonal relations (*β* = 0.644, *p* = 0.0045), which in turn related positively to receptive language at 24–26 months (*β* = 0.562, *p* = 0.0145) , whereas the direct *motor* → *speech* path was not retained (*β* = −0.453, *p* = 0.302). In the *social* → *motor* → *speech* example, early play leisure time related positively to later fine-motor skills (*β* = 0.483, *p* = 0.0018), and later fine-motor performance related positively to receptive language (*β* = 1.799, *p* = 0.0462), whereas the direct *social* → *speech* path was not retained (*β* = −0.784, *p* = 0.444). Full results for these exploratory motifs are summarized in Tables S9 and S10, and provided in the Supplementary Data 5 and 6.

Together, the cross-sectional and longitudinal follow-up analyses suggest that motor and social constructs may help organize distinct but related developmental routes toward later communication. Because the longitudinal analyses were based on modest complete-data subsets, these pathways are interpreted as hypothesis-generating rather than causal.

## 3 Discussion

This study was motivated by a central gap in the white-matter substrates of infant speech-acquisition (*1,11*). Although prior developmental neuroimaging work has yielded important insights into ventral and dorsal language pathways (*51, 52, 17*), most studies have focused either on a small set of canonical language tracts or on relatively restricted behavioral outcomes (*18*). As a result, it has remained unclear how language pathways co-develop with motor, auditory, and visual systems during the first two years of life, and whether early communication is better understood as a distributed developmental process rather than as the output of a single dominant tract. By combining longitudinal diffusion MRI, infant-oriented tract extraction across motor, auditory, visual, ventral-language, and dorsal-language systems, and a behavioral framework spanning motor, social interaction, visual/cognitive, daily living, emotion/regulation, and speech-related outcomes, the present study addresses that gap with a more explicitly multisystem developmental design.

A first major implication concerns the structural organization of tract maturation. Our data do not simply show that several tracts are “more mature” than others in an absolute sense. Rather, within the representative left-hemisphere tract set, the motor and auditory tracts occupied the upper end of the predicted near-birth NDI range, whereas AF, SLF II, and SLF III occupied the lower end. This matters because it situates dual-stream language pathways within a broader motor- and sensory-linked maturational landscape rather than treating them in isolation. At the same time, our high-angular-resolution diffusion imaging (HARDI) / constrained spherical deconvolution (CSD) based tractography detected sparse but plausible neonatal AF extension toward inferior frontal territory, while still confirming predominant terminations in precentral cortex. We do not interpret this as evidence for an adult-like language circuit at birth. Instead, it refines earlier lower-resolution conclusions (*48, 49*) by suggesting that a weak structural scaffold toward inferior frontal cortex may already be present (*23*), even though premotor connectivity remains the dominant early pattern. This interpretation is also consistent with the view that the dorsal pathway comprises an earlier-developing premotor component relevant to auditory–motor mapping and a later-maturing BA44 component more relevant to higher-order language operations (*Angela D. Friederici, personal communication, May 2025*).

The developmental course of these association tracts is equally important. AF, SLF II, SLF III, and CC_4_ showed closely aligned trajectory shapes and relatively steep log-age slopes; and by 24 months, AF, SLF II, and SLF III approached the maturity range of primary sensory tracts such as AR and OR. The refined interpretation of early white-matter maturation is therefore not simply one of rapid versus slow development, but one of progressive cross-system rebalancing: tracts begin from markedly different maturational states and move toward a more coordinated structural configuration over the first two years of life. Supplementary system-level analysis sharpened this interpretation. Inter-tract dispersion declined with age, indicating that the maturational landscape became progressively less heterogeneous (Figure 3 **b** and **d**, Figure S13), while motor-leading gradients anchored on CC_4_ were more informative than sensory–sensory contrasts, particularly for association tracts and predominantly motor behavioral outcomes. Together, these results suggest structured reorganization rather than diffuse convergence, with motor-linked architecture providing a major axis around which association pathways become coordinated.

A second major implication concerns the distributed and complex nature of tract–behavior couplings. Significant associations were not confined to canonical language tracts or to speech-related outcomes alone. Instead, they extended across motor, social interaction, visual/cognitive, daily living, emotion/regulation, and speech-related outcomes, and were distributed across motor, auditory, visual, ventral-language, and dorsal-language pathways. The same tract can relate to multiple developmental domains, and the same speech-related outcome can relate to multiple tract systems (*53*). This many-to-many pattern extends prior focused structure–function observations (*54, 55, 20, 18*) and supports an emergent account of early communication.

Within this distributed pattern, the motor domain was especially prominent. Motor outcomes showed broadly positive associations across the tract system, indicating that greater tract maturity generally covaried with stronger age-adjusted motor performance. Speech-related outcomes showed a more differentiated profile. Lexical measures were characterized by widespread predominantly negative linear age-adjusted associations, whereas gesture-related measures showed a sparser and partly motor-like pattern. These negative coefficients should not be read as evidence that greater white-matter maturation impairs communication. Rather, they are more consistent with timing-sensitive developmental coupling during a period in which precursor skills and later communication outcomes do not evolve synchronously (*56, 57*). The distinction between lexical and gesture measures further suggests that speech-related behavior is not a unitary infant construct. Gesture-based measures likely index earlier, more action-linked communicative stages, whereas lexical measures appear to depend on broader multisystem coordination involving motor, auditory, visual, and social development. This interpretation is consistent with treating speech-related outcomes as downstream expressions of a broader developmental cascade rather than as the only exclusive property of the system.

A third implication concerns the role of motor and social intermediates. The exploratory mediation analyses were not designed to establish a single definitive pathway. Instead, they asked whether the distributed tract–behavior associations could be statistically organized by intermediate developmental constructs. Here again, motor was central. In the cross-sectional screen, motor mediators strongly predominated among retained tract-to-speech triplets. The targeted longitudinal follow-up analyses, although based on modest complete-data subsets, further suggested *consistent candidates*: *motor* → *social* → *speech* and *social* → *motor* → *speech* motifs. These directions are not contradictory. Rather, taken together, they point to a broader developmental loop in which early action, social interaction, and later communication may be mutually reinforcing.

This interpretation resonates with developmental accounts that view gross motor skills as an early communicative resource in facilitating social and communication development during infancy (*58, 59, 12, 60, 61, 8*), social engagement as a catalyst for structured learning (*43, 62, 63, 64*), and fine-motor refinement as an increasingly important substrate for later cross-modal sensorimotor learning (*26, 65, 66, 23, 67, 13*). We therefore interpret these cascades as hypothesis-generating evidence for coordinated developmental organization, not as proof of a fixed causal sequence.

Placed in the context of the wider field, these results help bridge two traditions that have often remained partially disconnected. One tradition focuses on the anatomy of ventral and dorsal language tracts (*51, 52, 54, 55, 20, 18*). The other emphasizes sensorimotor association, social interaction, and developmental precursors in early learning (*60, 61, 66, 67, 68, 64, 8*). Our data suggest that these are not competing explanations. Rather, speech acquisition is better understood as an experience-dependent developmental process in which communication emerges through coordinated maturation across multiple neural systems and the precursor skills that link them (*6, 69, 27, 13*). Structurally, language-relevant association tracts develop within a broader field of motor, auditory, and visual pathways. Behaviorally, speech-related outcomes are embedded in multisystem patterns rather than mapped exclusively onto canonical language tracts. In this sense, our findings are relevant to debates about neonatal imitation and early sensorimotor readiness (*70, 71, 72, 73, 74*), because they support the presence of early sensory–motor structural substrates while also indicating that effective multisystem integration remains developmentally immature and likely depends on postnatal experience.

Although our structural data do not directly measure phonetic learning, one plausible functional correlate of this reorganization is experience-dependent perceptual attunement during the first post-natal year (*15, 75, 7*). This process is often described behaviorally as *perceptual narrowing* (*76*), but from a neurobiological perspective it is better understood as *perceptual reorganization*, encompassing both initial neural organization and experience-dependent change in perceptual sensitivity, including maintenance/loss as well as sharpening/improvement (*Janet F. Werker, personal communication, January 2025*). Our structural results extend, rather than overturn, previous reports that dorsal pathways are relatively immature near birth but change rapidly after birth (*77, 78, 52*). The key refinement here is that dorsal-language maturation is best understood relative not only to ventral-language tracts, but also to motor, auditory, and visual systems. Motor-leading motor-sensory gradients appear to consistently link to the maturation of AF, SLF II, and SLF III, as well as to the development of motor-skills (both gross- and fine-motor). This resonates with the prominent role of early motor experience in the sensorimotor foundation underlying perceptual reorganization (*23*).

Several limitations should be acknowledged. First, our dMRI results characterize structural microdevelopment rather than functional engagement; direct evidence of how active these circuits are at birth would require complementary measures such as EEG, MEG, fNIRS, or infant fMRI. Second, the behavioral battery indexes spoken communication and its developmental precursors rather than direct acoustic measures of speech production such as babbling rate. Third, many assessments rely on caregiver report or structured interview, raising the possibility of shared method variance across domains. Fourth, exact sample size varies across tract–behavior and mediation analyses. Fifth, the longitudinal mediation analyses were performed in modest complete-data subsets and were therefore best interpreted as exploratory. Finally, although supplementary tensor-based analyses were retained for benchmarking, the main inferences of the paper rest on NODDI-derived NDI because it provided more coherent and developmentally interpretable patterns than FA, MD, or RD in this infant dataset.

These limitations notwithstanding, the study has important strengths. It leverages one of the largest longitudinal infant HARDI resources currently available, uses infant-oriented tract extraction methods, explicitly broadens analysis beyond the classical ventral/dorsal language framing, and relates white-matter maturation to a multi-domain behavioral framework rather than to language outcomes alone. More importantly, it reframes the structural basis of infant speech acquisition as a question of coordinated multisystem development. The main contribution of this work is therefore not only to describe individual tract trajectories, but to show how motor, auditory, visual, ventral-language, and dorsal-language pathways become progressively organized into a more balanced developmental architecture that is behaviorally relevant for emerging communication.

Future work should move in two complementary directions. One is methodological: denser longitudinal sampling, multimodal imaging, and more direct speech-production phenotyping are needed to clarify how structural and functional development align across infancy. The other is conceptual: rather than asking whether a single pathway “causes” language, future studies should test how multiple functionally segregated but developmentally coordinated systems jointly scaffold the emergence of communication. Our results suggest that this broader, multisystem framework is not only theoretically preferable, but empirically necessary.

## 4 Concluding remarks

Across the first 24 months of life, infant white-matter maturation underlying speech acquisition is best understood as a coordinated multisystem process rather than as the isolated development of a canonical language pathway. Dorsal-language association tracts begin from relatively low predicted near-birth maturity yet undergo steep postnatal growth, progressively rebalancing with motor, auditory, and visual systems. This reorganization appears structured rather than diffuse, as suggested by declining inter-tract dispersion and the greater informativeness of motor-leading sensory–motor gradients than sensory–sensory gradients. The same multisystem pattern is also evident at the brain–behavior level: tract–behavior coupling is distributed across multiple tract systems and developmental domains, indicating that emerging communication is embedded in broad multisystem organization rather than mapped onto a single tract class or behavioral endpoint. Within this distributed landscape, motor and social capacities stand out as plausible organizing precursors, as suggested by the prominence of motor-linked associations and by exploratory follow-up cascades linking motor, social, and later speech-related outcomes. Rather than displacing dual-stream accounts of early language development, the present study situates them within a broader developmental framework in which motor, auditory, visual, ventral-language, and dorsal-language systems mature in concert to scaffold the emergence of communication. The central contribution of this work is therefore not simply to describe individual tract trajectories and tract–behavior associations, but to provide empirical support for understanding early speech acquisition as a problem of coordinated multisystem development.

## 5 Methods

### Participants, cohort derivation, and quality control

We analyzed infant neuroimaging and behavioral data from the Baby Connectome Project (BCP), a longitudinal cohort designed to characterize early brain and behavioral development (*45*). Analyses were restricted to the 0–24 month age window. Scan counting was performed at the visit level, such that multiple diffusion runs acquired within the same imaging visit were treated as a single scan session.

Cohort derivation proceeded in four stages (Figure 1; Table 2). First, the 0–24 month imaging inventory comprised 518 scan sessions from 234 infants. Second, 369 scan sessions from 197 infants had accessible imaging data and corresponding processing records. Third, after acquisition and processing quality control (QC), 255 scan sessions from 151 infants were retained in the processing-QC cohort. Finally, a representative tract-analysis sample comprised approximately 234 scan sessions from approximately 134 infants after tract-level plausibility checks and tract–behavior matching, with exact denominators varying modestly across tract- and behavior-specific analyses. QC decisions were based on predefined imaging and processing criteria and were applied independently of behavioral outcomes. Exclusion criteria included: (*i*) acquisition issues, such as early termination, severe motion, blurring, and signal loss; (*ii*) processing failures, including distortion-correction failure, severe intensity artifacts, or misalignment; and (*iii*) tract extraction failures or implausible longitudinal outliers in individualized tract maturation trajectories. At the diffusion-run level, a scan session was retained for processing-QC if at least one diffusion run was rated as acceptable (pass or questionable) after eddy-correction QC. Demographic and sampling characteristics of the processing-QC cohort are summarized in Table 2.

### MRI acquisition and data processing

The BCP acquisition protocol has been described previously (*45*). MRI was acquired on a 3.0T Siemens Prisma scanner. T1w data were obtained using a magnetization-prepared rapid gradient-echo (MPRAGE) sequence with the following parameters: TR = 2400 ms; TE = 2.24 ms; flip angle = 8^◦^, matrix size = 320 × 320, field of view (FOV) = 256 × 256 mm^2^; and isotropic voxel size of 0.8 mm^3^.

The dMRI data were acquired using a multiband spin-echo echo-planar imaging (EPI) sequence with the following parameters: TR = 2640 ms; TE = 88.6 ms; flip angle = 78^◦^, matrix size = 140×105, FOV = 210 × 210 mm^2^; and isotropic voxel size = 1.5 mm^3^. The diffusion acquisition included 144 diffusion-weighted directions distributed across 6 shells (*b* = 500, 1000, 1500, 2000, 2500, and 3000 s/mm^2^), together with 6 *b*_0_ volumes. To ensure longitudinal compatibility with the HCP Lifespan Pilot Project (*79*), two specific shells were explicitly incorporated.

Structural and diffusion MRI data were processed using an infant-oriented multimodal pipeline built on FSL (*80*), ANTs (*81*), and MRtrix3 (*82*), together with our previously developed infant processing tools (*46, 21, 22*). Processing steps included brain extraction, infant-oriented tissue segmentation, cross-modal registration between sMRI and dMRI, estimation of fiber orientation distributions, and tractography in native diffusion space. These procedures were designed to accommodate the marked anatomical and tissue-contrast variability that characterizes the first two postnatal years. Additional anatomical validation of the tractography framework is provided in S2. Additional tract extraction and anatomical validation.

### Tract extraction and tract systems

White-matter tract extraction was performed using an extension of our previously developed NAT framework (*22*), using protocols derived from the Human Connectome Project (HCP) dataset (*83, 84, 85*). Rather than imposing adult tract geometry directly onto infant brains, the adult atlas served as a source of tract definitions together with endpoint and inclusion–exclusion ROI priors. Atlas-to-infant alignment was driven by infant-specific tissue segmentation and anatomical registration, after which tractography was conducted in each infant’s native dMRI space.

Tractography used multi-shell, multi-tissue constrained spherical deconvolution (CSD) to estimate fiber orientation distributions (*86*), together with anatomically constrained tractography (ACT) to enforce biologically plausible streamline propagation and termination (*82*). Additional infant-oriented strategies included adaptive thresholding and ROI dilation to improve robustness in immature tissue while limiting spurious streamlines. Post-registration overlays of tissue maps and tract endpoint ROIs were visually inspected, and scans with severe segmentation or registration failures were excluded.

Across the full study framework, 33 tracts were organized into five systems: auditory, visual, motor, ventral-language, and dorsal-language (Figure 2), with each system represented consistently by a distinct color in the text and legend. The auditory system comprised peripheral, subcortical, and cortical auditory tracts; peripheral auditory tracts were excluded from tract-level microstructural analyses because of excessive measurement noise. The visual system included the optic radiation (OR), middle longitudinal fasciculus (MLF), and SLF II. Although MLF and SLF II are sometimes discussed within broader language frameworks (*11*), we grouped them with the visual system because of their established roles in auditory–visual–semantic integration and visual–motor integration (*87, 88, 89*). The motor system included the corticospinal tract (CST), thalamo-precentral tract (T-PREC), and CC_4_. The ventral-language system comprised the UF, inferior fronto-occipital fasciculus (IFOF), and ILF, with IFOF highlighted separately in figures to reflect its dual relevance to visual and semantic processing (*90*). The dorsal-language system comprised the AF and SLF III (*91*).

For compact developmental visualization, main-text trajectory and maturity-structure summaries used a representative left-hemisphere tract set spanning these systems. Tract–behavior analyses used the full 33-tract framework.

### NODDI modeling and tract-level NDI summaries

We used NODDI to estimate tract microstructure from the multi-shell diffusion data (*47*). The primary metric used throughout the study was the neurite density index (NDI), interpreted as an index of intra-neurite volume fraction and neurite packing density. We selected NDI as the primary microstructural outcome because it provides a biologically motivated compartment-based measure that is more appropriate than single-tensor metrics for multi-shell data and for regions with complex fiber geometry.

Voxel-wise NDI maps were computed and then summarized at the tract level by averaging NDI values within each tract mask in native diffusion space. These mean tract NDI values formed the basis for developmental trajectory analyses, tract–behavior association models, and exploratory mediation analyses. Tensor-derived metrics including FA, MD, and RD were evaluated as supplementary benchmarks but were not used as primary inferential measures.

### Behavioral assessments and domain framework

Behavioral variables were derived from 11 BCP assessments spanning six broad developmental domains: motor, social interaction, visual/cognitive, daily living, emotion/regulation, and speech-related outcomes. These assessments included parent-report questionnaires, structured caregiver interviews, and examiner-administered tasks. Behavioral scores were matched to diffusion MRI visits using a temporal window of ±1 month.

These 11 behavioral assessments were categorized into two groups — *Primary* and *Supple-mentary* — as summarized in Table 1. The *Primary* group comprised 5 assessments used across all main analytical stages: tract–behavior correlation analysis, cross-sectional mediation screening, and longitudinal exploratory follow-up. Each primary assessment provided more than 70 matched observations for tract–behavior analyses. The *Supplementary* group included 6 additional assessments that were used only in tract–behavior correlation analyses because of limited matched sample sizes. Model-specific matched tract–behavior sample counts are provided in the Supplementary Data 1.

For tract–behavior screening, we retained variables with sufficient visit-level coverage after MRI matching. The full instrument list, administration mode, age range, domain assignment, and analytical usage are summarized in Table 1. For visualization in the tract–behavior heatmap, speech-related outcomes were further divided into lexical measures and gesture-related measures, yielding seven displayed heatmap columns from an underlying six-domain behavioral framework.

### Statistical analysis

For infant *i*, visit *j* , and tract *t*, let 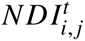 denote the mean NDI of a tract, *Age*_*i*, *j*_ denote postnatal age in months, and *Sex*_*i*_ denote sex. Subject-specific random effects are denoted by *u*_*i*_, and residual errors by 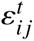. Fixed-effect population-level predictions for tract *t* at age *Age* are denoted by 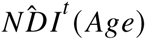 Unless otherwise noted, age is measured in postnatal months.

#### Single-tract developmental trajectory models

To characterize population-level tract maturation across infancy, we fitted tract-wise mixed-effects models to mean tract NDI values as a function of postnatal age (*92*). For descriptive trajectory visualization, age was modeled with a quadratic fixed-effect function to allow smoothly decelerating maturation over the 0–24 month interval:

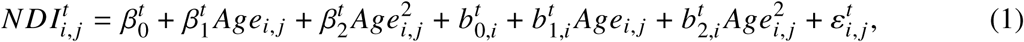

equivalently, in Wilkinson notation,

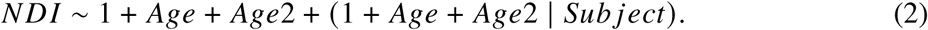

where, *Age*2 = *Age*^2^, denoting the squared age term. Candidate age functions included linear, logarithmic, and quadratic specifications, which are commonly used in developmental neuroimaging research (*93,94,20*); model comparisons are reported in the Tables S3, S4, and S5. For developmental visualization, we used a common quadratic age specification, selected to provide physiologically plausible and population-level predicted trajectories across tracts (Figures 3 **a**, S5, S6, S7, S8, S9, S10, S11, and S12).

Because each infant contributed a limited number of repeated scans (Table 2), these models were used to primarily estimate population-level developmental trajectories rather than subject-specific quadratic growth curves. To directly compare between-tract maturation-rate parameter, we instead used single-tract log-age mixed-effects models for maturation-rate estimation:

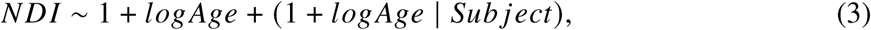

where *logAge* = log(1 + *Age*). The tract-specific fixed-effect coefficient for *logAge* was taken as the absolute maturation-rate estimate reported in Table 3.

#### Measures for quantitative analysis across trajectories

To quantify trajectory-shape similarity, the fitted tract curves were evaluated on the age grid *τ* ∈ [0, 24]. For each tract pair (*r*, *c*), we first computed the age-wise difference,

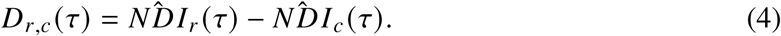

We then defined the shape-distance between the two curves as the standard deviation of this difference across age:

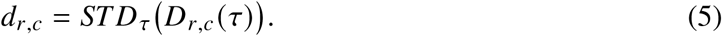

Finally, we mapped this distance to a bounded similarity score,

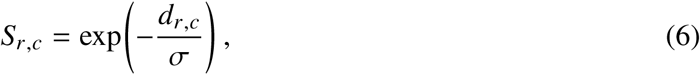

where *σ* was set to the median non-zero pairwise shape-distance across tract pairs. Similarity values closer to 1 indicate more parallel trajectory shapes, whereas lower values indicate greater divergence across age, as shown in Figure 3 **c**.

To compare developmental slopes across tracts within a unified framework, we fitted a single mixed-effects model jointly across tracts,

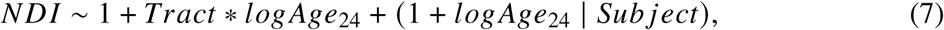

where *logAge*_24_ = log(1+*Age*)−log(1+24). Centering this term at 24 months sets *logAge*_24_ = 0 at the late-infancy endpoint; thus, tract main effects reflect predicted maturity differences at 24 months, whereas tract-by-logAge_24_ interactions reflect differences in log-age maturation rate relative to the categorical reference tract. In the implementation, AF_*L*_ was used as the categorical reference; this choice only defines coefficient coding, because all reported pairwise slope contrasts and maturity-difference matrices were obtained as linear contrasts or fixed-effect predictions from the same joint model. For Figure 3 **c**, *p*-values from the 136 pairwise slope contrasts among the 17 displayed representative tracts were adjusted using BH–FDR; markers denote tract pairs whose slope difference was not significant after correction (*q* ≥ 0.05). These markers indicate absence of evidence for different slopes within the fitted model, rather than formal equivalence.

We used this model in two ways. First, an omnibus test of the tract-by-logAge_24_ interaction evaluated whether developmental slopes differed across tracts overall, and pairwise comparisons from this model were used to identify tract pairs whose slopes did not significantly differ, as denoted by the overlaid markers in the similarity matrix of Figure 3 **c**. Second, fixed-effect predictions from the same joint model were used to construct pairwise maturity-difference matrices at 2 weeks and 24 months, as shown in Figure 3 **b** and **d**.

For a given evaluation age *Age*, we defined the signed pairwise maturity difference between tracts *r* and *c* as

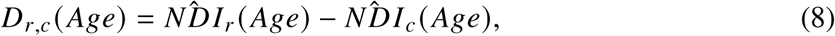

where 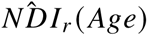 denotes the fixed-effect predicted NDI of tract *r* at age *Age*. Thus, each matrix entry represents the predicted NDI of the row tract minus that of the column tract (Figure 3 **b** and **d**) . Values near 0 indicate more similar maturity levels, positive values indicate that the row tract is higher, and negative values indicate that the row tract is lower. Near-birth matrices were evaluated at *Age* = 0.46 months (approximately 2 weeks), and late-infancy matrices at *Age* = 24 months.

#### Tract–behavior association models

For each tract–behavior pair, we fitted an age- and sex-adjusted mixed-effects model with a subject-specific random intercept. Behavioral score served as the dependent variable, and tract-level NDI entered as both linear and quadratic fixed effects:

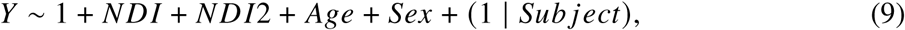

where *NDI*2 = *NDI*^2^, denoting the squared tract-level NDI term. This specification allowed both approximately monotonic and curvilinear tract–behavior relationships while accounting for repeated visits and developmental timing. A tract–behavior association was retained if the linear NDI term survived BH–FDR correction across 33 tracts within that behavioral variable. The domain-level heatmap in Figure 4 summarizes the number of significant tract–behavior associations, and the sign displayed within each tract-by-domain cell corresponds linear NDI component and is therefore used only as a directional summary, not as a complete description of curvilinear effects.

#### Cross-sectional mediation screening

To organize distributed tract–behavior associations into candidate developmental pathways, we performed an exploratory cross-sectional mediation screen using piecewise structural equation models (SEMs) implemented in R via piecewiseSEM (*95*). Triplets took the form predictor (*X*), mediator (*M*), and outcome (*Y*), where *X* could be either a tract-level NDI variable or a behavioral measure. Candidate mediators and outcomes were selected from behavioral variables spanning the six developmental domains according to 19 prespecified pathway families summarized in Table S8, with candidate variables listed in Table S2. Exact testable and retained triplet counts are reported in the Results.

For each triplet ( *X*, *M*, *Y*), we fitted paired mediator and outcome models:

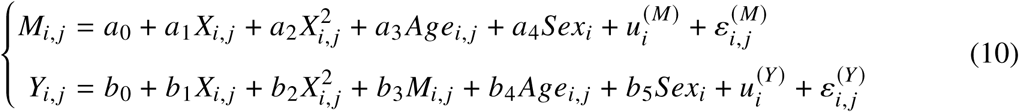

Quadratic terms in *X* were included when tract-level NDI served as the predictor, allowing non-linear tract effects. Age and sex were included as covariates, and repeated visits were modeled using subject-specific random intercepts.

Because behavioral and neural variables were not available at all visits, missing values were handled using predictive mean matching (PMM) in the mice package of R, generating 100 imputed datasets (*96*). Model parameters were estimated separately within each imputed dataset and subsequently pooled using Rubin’s rules (*97*), effectively accounting for both within- and between-imputation variability. Pooled *p*-values were calculated from the corresponding *t*-statistics, using the combined standard errors and degrees of freedom under Rubin’s combination formula.

To retain a triplet, both the *X* → *M* path and the *M* → *Y* path controlling for *X* were required to survive a study-wide BH–FDR correction across all testable triplets (*q* < 0.05). These cross-sectional mediation results were interpreted as prioritization and hypothesis generation rather than as evidence of causal mediation.

#### Longitudinal exploratory follow-up

Motivated by the cross-sectional enrichment of motor and social intermediate constructs, we conducted targeted time-ordered follow-up analyses using matched complete-data behavioral observations without imputation. We tested triplets of the form predictor at approximately 6 months (4–6 months), mediator at approximately 12 months (12–14 months), and outcome at approximately 24 months (24–26 months):

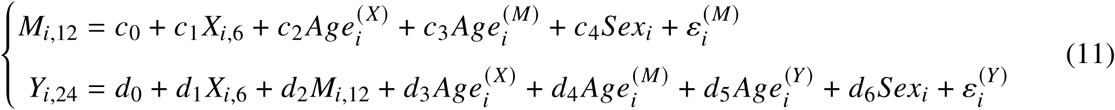

where *Age_*i*_* ^(*X*)^, *Age_*i*_* ^(*M*)^, and *Age_*i*_* ^(*Y*)^ denote the exact ages within the predictor, mediator, and outcome windows, respectively.

Two developmental motifs were examined in modest complete-data subsets (*N* ≈ 35–45): (*i*) *motor* → *social* → *speech*, and (*ii*) *social* → *motor* → *speech*. For each motif, we used the piecewise SEM framework (*95*) with two linear regression models following (*67*). The mediator was predicted from the earlier predictor, and the later outcome was predicted from both the earlier predictor and the later mediator. Exact ages within the sampling windows and sex were included as covariates when available.

Because these longitudinal analyses were targeted exploratory follow-up analyses in modest complete-data subsets, we did not apply BH–FDR correction (*50*). We reported uncorrected *p* values, effect estimates, and uncertainty, and interpret these analyses as hypothesis generating rather than causal.

#### Multiple-comparison correction

Unless otherwise specified, multiplicity was controlled using the BH–FDR. For tract–behavior screening, correction was applied across the 33 tracts within each behavioral variable. For cross-sectional mediation screening, correction was applied study-wide across all 74,276 testable triplets. The longitudinal follow-up analyses were targeted and exploratory; accordingly, we reported uncorrected *p*-values and interpret these results cautiously.

## Author contributions

F.L. designed the research, wrote the code, and drafted the manuscript. F.L., J.L., and J.G. collected and processed the dataset. F.L. and X.C. provided statistical analysis and interpretation of the data. F.L., Y.W., Z.W., and H.W. made contributions in the areas of anatomical delineation and data analysis. F.L., J.F., H.Z., and D.S. coordinated and supervised the whole work. All authors were involved in critical revisions of the manuscript, and have read and approved the final version.

## Additional acknowledgments

We gratefully acknowledge Professors *Janet F. Werker* and *Angela D. Friederici* for their invaluable insights and constructive suggestions provided through personal communications, which greatly enhanced our interpretation and presentation of the data. Specifically, Professor Janet F. Werker provided critical clarifications on terminology and conceptual distinctions among perceptual attunement, perceptual narrowing, and perceptual reorganization (*January 2025*). Professor Angela D. Friederici clarified the anatomical and functional differentiation within dorsal language pathways, highlighting distinct roles of pathways to premotor cortex (BA 6) and inferior frontal cortex (BA 44) (*May 2025*). Their contributions significantly improved the clarity and scholarly depth of our manuscript.

## Data and materials availability

Raw BCP MRI and behavioral data are available through the Lifespan Baby Connectome Project / Human Connectome Project portal —. https://www.humanconnectome.org/study/lifespan-baby-connectome-project. Source data supporting Figures 3–5 and the reported tract–behavior and pathway analyses are provided as Supplementary Data 1–6. Image processing and tractography were performed using FSL, ANTs, MRtrix3, and our infant-oriented processing tools (*80, 81, 82, 46, 21, 22*). Statistical analyses were implemented in MATLAB and R. Figure generation combined MATLAB- and TikZ-based workflows. Publicly available resources associated with our tract-processing framework are provided in the cited software references.

## Supplementary materials

Supplementary Methods

Supplementary Results

Figures S1 to S14

Tables S1 to S12

References (*108–116*)

## Supplementary files

Supplementary Data 1: Counts of matched tract–behavior samples, Supplementary Data 2: Significant tract–behavior associations ordered by behavior, Supplementary Data 3: Significant tract–behavior associations ordered by tract, Supplementary Data 4: Cross-sectional pathway-prioritization results, Supplementary Data 5: Longitudinal *motor* → *social* → *speech* results, Supplementary Data 6: Longitudinal *social* → *motor* → *speech* results.

