## Supplementary materials for "Diffusion Neuroimaging of Speech Acquisition in Infants"

### Supplementary Materials for
### Diffusion Neuroimaging of Speech Acquisition in Infants

Feihong Liu[1,2*], Jiameng Liu[1], Jinchen Gu[1], Yaoxuan Wang[3,4,5], Xinyi Cai[1], Zhaoyan Wang[3,4,5], Hao Wu[3,4,5], Jun Feng[2*], Han Zhang[1,6*], Dinggang Shen[1,6,7*]

**This PDF file includes:**

Supplementary Methods

Supplementary Results

Figures S1 to S14

Tables S1 to S12

**Supplementary files:**

The following supplementary files provide detailed tract-level and behavior-level statistical results:

`Supplementary Data 1`: Counts of matched tract–behavior samples,

`Supplementary Data 2`: Significant tract–behavior associations ordered by behavior,

`Supplementary Data 3`: Significant tract–behavior associations ordered by tract,

`Supplementary Data 4`: Cross-sectional pathway-prioritization results,

`Supplementary Data 5`: Longitudinal $motor \rightarrow social \rightarrow speech$ results,

`Supplementary Data 6`: Longitudinal $social \rightarrow motor \rightarrow speech$ results.

#### Supplementary Methods

#### S1. Additional cohort derivation and visit-level matching

We used data from the Baby Connectome Project (BCP), a large longitudinal cohort of infant neuroimaging and behavioral assessments (*45*). Within the 0–24 month diffusion MRI (dMRI) inventory, 518 scan sessions from 234 infants were identified. Session-level availability, processing quality control (QC), and tract-analysis QC were harmonized using a visit-based counting convention, such that one scan session corresponded to one imaging visit regardless of whether one or both diffusion runs were usable at that visit. After restricting to sessions with accessible diffusion data, 369 scan sessions from 197 infants remained. Following acquisition- and processing-level QC, 255 scan sessions from 151 infants were retained. The age distribution of retained sessions is summarized in Table S1. A further tract-analysis QC step yielded a representative tract-analysis sample of 234 scan sessions from 134 infants, with exact denominators varying modestly across tracts and behavior-specific models because tract extraction success and matched behavioral availability differed slightly by analysis.

QC decisions were based on predefined imaging and processing criteria rather than behavioral outcomes. These criteria included acquisition failure or severe motion, distortion-correction failure, gross misregistration, segmentation or tract-extraction failure, and tract measurements deemed implausible with respect to within-subject longitudinal profiles. Behavioral variables were matched to diffusion sessions within a ±1 month window. Consequently, model-specific sample sizes differed across behavioral outcomes and tracts and were generally smaller than the full imaging cohort. Model-specific matched counts are provided in the `Supplementary Data 1`, and representative sample-size contrasts across CDI-WG and MSEL are illustrated in Figure S1. For each behavioral assessment, the numbers of visits ($N_{\mathrm{obs}}$) and infants ($N_{\mathrm{sub}}$) are summarized in Table 1.

#### S2. Additional tract extraction and anatomical validation

Infant structural MRI (sMRI) and dMRI data were processed using our previously developed multimodal MRI pipeline (*46*), which integrates FSL (*80*), ANTs (*110*), and MRtrix3 (*82*). Brain tissue segmentation and region-of-interest (ROI) parcellation were performed using our deeplearning-based tools, optimized for longitudinal consistency across early development (*111*,*112*,*21*). These steps enabled accurate cross-modal registration between sMRI and dMRI (*113*), estimation of fiber orientation distributions (*86*), and tractography in infant native space (*114*).

Building on our NeoAudi Tract (NAT) toolbox (*85*), which was originally developed for individualized auditory-tract extraction from infant dMRI data, we extended the framework to additional white-matter systems. In brief, tract extraction proceeded in four stages:

1. HARDI data from 105 adult HCP subjects were used to construct an initial tract atlas (*83*,*115*).

2. A coarse atlas of target tracts was generated by averaging anatomically aligned adult tracts (*81*), following the previously validated auditory-tract strategy (*85*).

3. The atlas was iteratively refined through repeated extraction and alignment to yield a higher-resolution adult reference atlas.

4. The refined atlas was then transferred to individual infant dMRI data using the dual-branch registration framework of NAT, which was designed to preserve both cortical and subcortical alignment across infants of different ages.

By this strategy, cortical, subcortical, and peripheral ROIs were transferred using individualized tissue segmentation and brainstem parcellation rather than direct adult-to-infant intensity matching alone. Tractography was subsequently performed in native diffusion space using multi-shell multi-tissue constrained spherical deconvolution (CSD) and anatomically constrained tractography (ACT), with infant-tailored thresholds and ROI dilation to improve robustness while limiting anatomically implausible streamlines. Representative overlays of warped ROIs, tissue segmentations, and tract streamlines in infant native space were visually inspected as part of anatomical validation.

#### S3. Behavioral measure dictionary

For the primary statistical analyses, we selected 5 behavioral assessments that yielded larger numbers of matched tract–behavior pairs than the remaining 6 assessments. Behavioral variables used in tract–behavior screening and mediation analyses are summarized in Table S2, which serves as a merged variable dictionary. Measures were organized into six developmental domains: motor, social interaction, visual/cognitive, daily living, emotion/regulation, and speech-related outcomes.

The speech-related domain included vocabulary and phrase measures from CDI-WG, receptive and expressive language measures from the Mullen Scales of Early Learning (MSEL), and communication measures from the Vineland Adaptive Behavior Scales. Because the CDI-WG gesture-related phrase measure showed a tract-association pattern distinct from the other speech-related variables, it was displayed separately in the main tract–behavior heatmap (Figure 4), although conceptually it remains part of the broader speech-related outcome.

#### S4. Additional statistical models

##### S4.1. Model comparison for developmental trajectory charting

To visualize developmental trajectories, we evaluated tract-wise mixed-effects models with age as the primary predictor. Candidate age functions included linear, logarithmic, and quadratic specifications, which are commonly used in developmental neuroimaging research (*93, 94, 20*). Model-comparison results are reported in Tables S3–S5. These comparisons were used to justify the quadratic models employed for trajectory visualization in the main text, whereas log-age models were used separately for tract-wise maturation-rate estimation.

##### S4.2. Piecewise early-versus-late slope analyses

As a supporting robustness analysis, we additionally fitted piecewise mixed-effects models with a knot at 12 months to test whether tract maturation was steeper during the first postnatal year than during the second. These models yielded separate fixed-effect slope estimates for the 0–12 month and 12–24 month ranges. Across the representative left-hemisphere tract set, early-life slopes exceeded later-life slopes, supporting an interpretation that microstructural maturation was faster in the first postnatal year (Figure S2).

##### S4.3. Tensor-based benchmark analyses

As benchmark analyses, we applied the same repeated-measures modeling framework to tensor-derived diffusion metrics, including radial diffusivity (RD), mean diffusivity (MD), and fractional anisotropy (FA), to assess whether the broad developmental patterns observed for NDI were reproduced by less compartment-specific measures. These analyses were intended to provide context and robustness checks rather than to replace the primary compartment-based NODDI analysis.

##### S4.4. Raw-versus-standardized behavioral sensitivity analyses

Where both raw and age-normed standardized behavioral scores were available, both representations were retained for screening and later compared in a dedicated sensitivity analysis (Figure S3). This analysis was designed to evaluate whether tract-wise effect profiles depended on score scaling. Across paired raw and standardized outcomes, effect directions and tract-wise profiles were highly concordant, indicating that the qualitative tract–speech patterns are not driven by the choice of behavioral scaling.

### Supplementary Results

#### R1. Additional tract visualization

Figure S4 provides in-brain anatomical views of the neonatal left AF. Consistent with the main-text renderings, the neonatal AF terminated predominantly in precentral regions while also showing sparse but plausible extension toward inferior frontal territory, including pars opercularis and pars triangularis. This pattern is broadly consistent with previous reports emphasizing predominant neonatal precentral connectivity of the AF (*48*, *49*, *116*), while also illustrating the increased sensitivity of HARDI/CSD-based infant tractography to sparse inferior frontal extensions.

#### R2. Model comparison and robustness of trajectory charting

Tables S3, S4, and S5 summarize the model-comparison results for tract-wise developmental charting. Across the tract sets, quadratic models generally provided the best compromise between goodness of fit and physiologically plausible trajectory shape over the 0–24 month interval. Accordingly, quadratic models were used for developmental curve visualization, whereas separate log-age models were used for tract-wise absolute maturation-rate estimation.

Trajectories in Figures S5, S7, S9, and S10 and point plots in Figures S6, S8, S11, S12 provide additional context for both NDI and tensor-based benchmark metrics. The benchmark analyses using FA, MD, and RD reproduced previous findings about the relative maturity level between dorsal and ventral language pathways (*77, 78, 52*), except for left-hemisphere FA, where the AF even showed higher maturity than the UF at birth. However, they were less coherent than NDI overall. In particular, FA suggested implausibly low relative maturity for several auditory brainstem tracts compared with the AR, and tensor-derived metrics did not yield consistent associations with speech-related outcomes across CDI-WG, MSEL, and Vineland measures, as well as across RD, MD, and FA. For this reason, tensor-based analyses are presented as supporting benchmarks rather than as the primary basis for inference.

#### R3. Supporting system-level balance analyses

Figure S13 extends the tract-wise results in the main text to a system-level perspective. Inter-tract dispersion declined with age, indicating that the tract maturity landscape became progressively less heterogeneous over infancy. To quantify this pattern, we defined visit-wise inter-tract dispersion for infant $i$ at visit $j$ as the coefficient of variation of tract-level NDI across the representative tract set:

$$CV_{i,j} = \frac{STD\left(NDI_{i,j}^{t}\right)}{Mean\left(NDI_{i,j}^{t}\right)}, \tag{S1}$$

where $t$ denotes tracts. Lower $CV_{i,j}$ indicates a less heterogeneous tract-maturity profile. Age-related change in inter-tract dispersion was assessed using

$$CV \sim 1 + Age + Age2 + (1 + Age + Age2 \mid Subject). \tag{S2}$$

To examine balance between motor and sensory systems, we additionally defined gradient metrics using $CC_4$ as a motor anchor and AR/OR as sensory anchors:

$$\begin{cases} \nabla_{i,j}^{\text{M-AR}} = NDI_{i,j}^{CC_4} - NDI_{i,j}^{AR} \\ \nabla_{i,j}^{\text{M-OR}} = NDI_{i,j}^{CC_4} - NDI_{i,j}^{OR} \\ \nabla_{i,j}^{\text{AR-OR}} = NDI_{i,j}^{AR} - NDI_{i,j}^{OR} \end{cases} \tag{S3}$$

Associations of each gradient metric with outcome $Y$ (either an association-tract NDI or a behavioral measure) were tested using age- and sex-adjusted mixed-effects models with subject-specific random intercepts:

$$Y \sim \nabla + \nabla 2 + Age + Sex + (1 \mid Subject). \tag{S4}$$

where $\nabla$ denotes one of the gradient metrics of Eqn. S3, and $\nabla 2 = \nabla^2$.

These analyses indicated that the observed convergence was structured rather than diffuse. Motor-leading contrasts defined relative to $CC_4$ showed substantially more significant associations with association-tract maturity than did sensory–sensory contrasts between OR and AR. Specifically, the motor-leading gradients yielded 14, 13, 11, and 8 significant association-tract outcomes, whereas the OR–AR gradients yielded only 2 and 0 such outcomes (Table S6). Behavioral associations were fewer and were concentrated mainly in motor outcomes, with lower-$N$ CBCL- and ITSEA-derived associations interpreted as exploratory because of limited precision and coverage (Table S7). Together, these findings support the interpretation that progressive rebalancing across infancy is organized partly around a motor-linked axis rather than reflecting an undifferentiated flattening of tract differences.

#### R4. Additional tract–behavior displays and sensitivity analyses

Figure S14 provides additional representative tract–behavior examples beyond those shown in the main text. These plots illustrate the range of positive, negative, and curvilinear associations summarized at the domain level in the main tract–behavior heatmap (Figure 4). Each point denotes one visit, and thin gray lines connect repeated visits within infant. Black curves and shaded bands show fixed-effect predictions from the corresponding age- and sex-adjusted mixed-effects model, with subject-specific random effects set to zero. These curves therefore represent population-level adjusted associations rather than nonparametric smoothers fit directly to the raw data.

Because raw and age-normed standardized versions of some behavioral measures can differ substantially in scale, we additionally evaluated whether tract–behavior results depended on behavioral scaling. Figure S3 compares tract-wise effect-size profiles obtained from raw versus age-normed standardized outcomes. Across matched constructs, directions and tract-wise profiles were generally concordant, indicating that the main tract–behavior interpretation was not driven by a particular score-scaling choice.

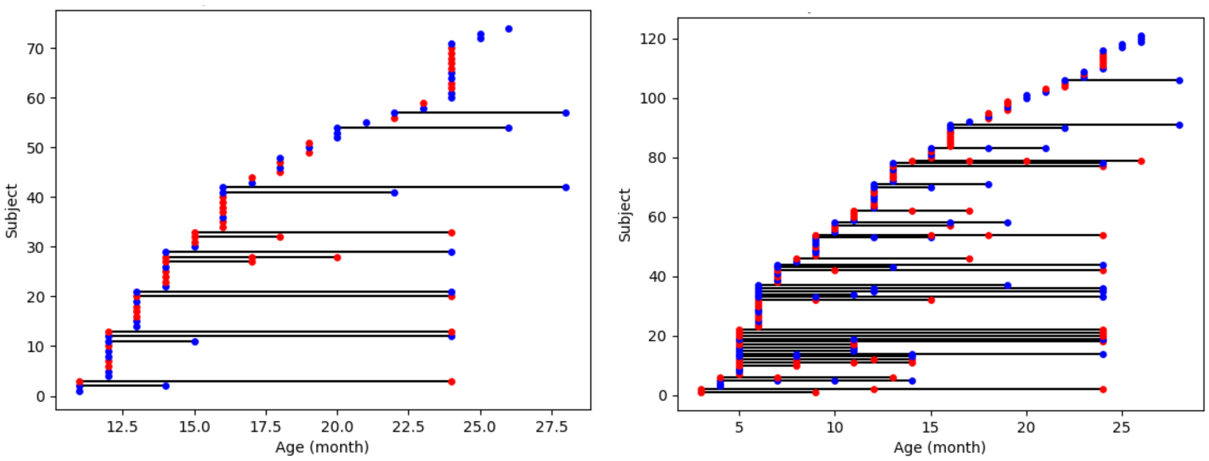


**Figure S1**: **Model-specific matched sample sizes and visit distributions for representative speech-related instruments. Left**, CDI-WG; **right**, MSEL. Each point denotes one visit-level behavioral observation matched to dMRI within the prespecified temporal window; points are colored by infant sex, and horizontal black lines connect repeated visits within infant. This figure illustrates why model-specific tract–behavior sample sizes are smaller than the full imaging and behavioral assessment cohort and why matched coverage varies across instruments.

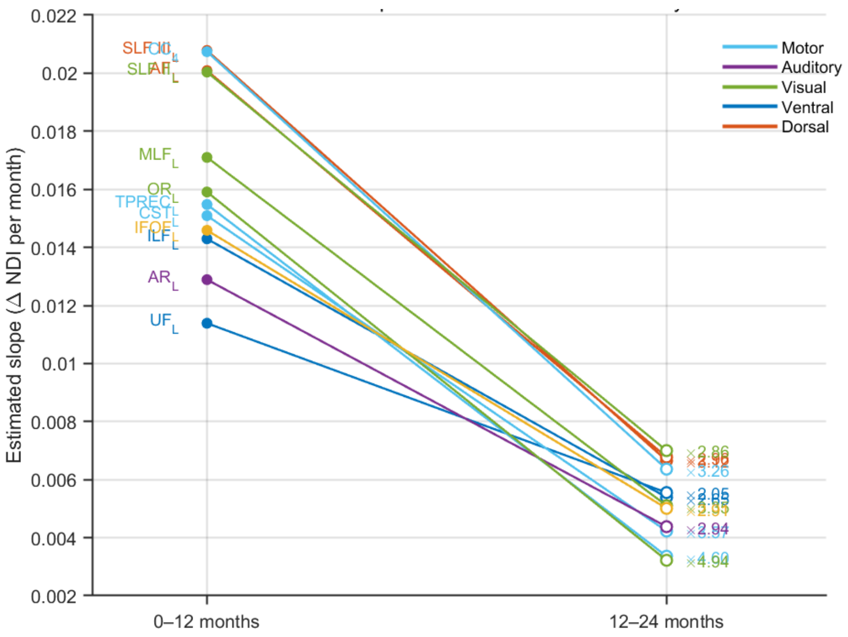


**Figure S2**: **Piecewise NDI slope estimates across the first and second postnatal years.** Each colored line connects the estimated monthly NDI slope for a given representative tract in the 0–12-month window and the corresponding slope in the 12–24-month window, derived from piecewise mixed-effects models. Colors denote tract systems. Labels on the left identify tracts, and the colored multipliers on the right indicate the ratio of the first-year slope to the second-year slope for each tract. Across tracts, estimated NDI slopes were consistently larger in the first year than in the second, indicating steeper early postnatal maturation.

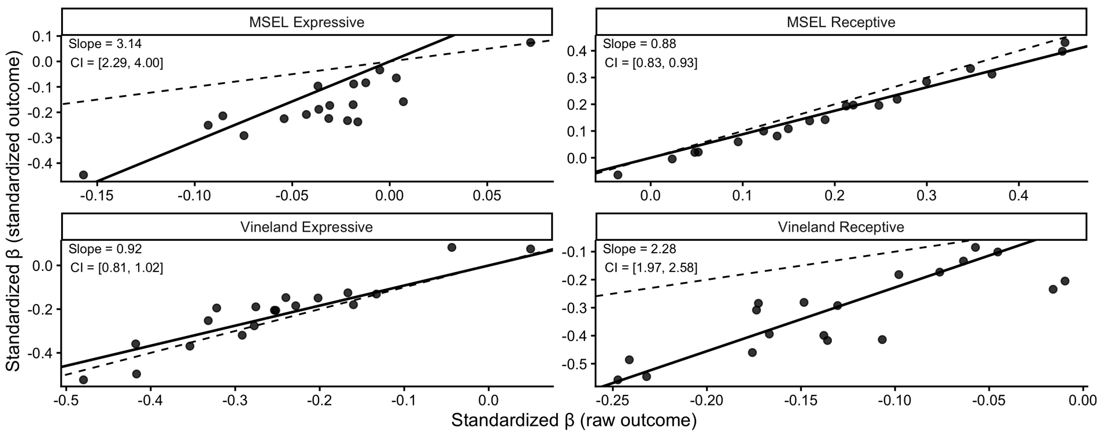


**Figure S3**: **Concordance of tract-wise effect-size profiles using raw versus age-normed standardized speech-related measures.** Each panel shows one matched raw/standardized speech construct. Each point represents one tract; the $x$-axis shows the standardized fixed-effect coefficient from the age- and sex-adjusted mixed-effects model using the raw score, and the $y$-axis shows the corresponding coefficient from the matched model using the age-normed standardized score. Both models used the same specification, $Outcome \sim NDI + NDI2 + Age + Sex + (1 \mid Subject)$. The dashed line indicates equality ($y = x$).

**Table S1**: **Age distribution of retained diffusion MRI sessions after acquisition and processing QC.** Entries report the number of retained scan sessions at each age bin between 2 weeks and 24 months under the visit-based counting convention used throughout the study.

| **Age** | **2 wk** | **1 mo** | **2 mo** | **3 mo** | **4 mo** | **5 mo** | **6 mo** | **7 mo** | **8 mo** | **9 mo** | **10 mo** | **11 mo** | **12 mo** |
|---|---|---|---|---|---|---|---|---|---|---|---|---|---|
| Sample size ($N$) | 5 | 2 | 1 | 9 | 12 | 17 | 15 | 10 | 15 | 17 | 7 | 13 | 17 |
| **Age** | | **13 mo** | **14 mo** | **15 mo** | **16 mo** | **17 mo** | **18 mo** | **19 mo** | **20 mo** | **21 mo** | **22 mo** | **23 mo** | **24 mo** |
| Sample size ($N$) | | 10 | 11 | 15 | 10 | 5 | 8 | 8 | 6 | 5 | 6 | 6 | 25 |

**Table S2**: **Merged behavioral variable dictionary across the six developmental domains used in tract–behavior and mediation analyses.** The table lists each behavioral measure/score and its source instrument. Speech-related measures are listed under a single domain here for analytical completeness, although one gesture-related measure was displayed separately in the main-text heatmap (Figure 4) to preserve interpretability.

| **Measure / score** | **Source instrument** |
|---|---|
| **Domain 1: Motor** | |
| Early gestures total | CDI-WG |
| Later gestures total | CDI-WG |
| Total gestures total | CDI-WG |
| Gross Motor T Score | MSEL |
| Fine Motor T Score | MSEL |
| Gross Motor Raw Score | MSEL |
| Fine Motor Raw Score | MSEL |
| Gross subdomain Raw Score | Vineland |
| Fine subdomain Raw Score | Vineland |

*Continued on next page*

*Table S2 continued*

| Measure / score | Source instrument |
|---|---|
| Gross subdomain V-scale Score | Vineland |
| Fine subdomain V-scale Score | Vineland |
| Motor skills domain V-scale Score | Vineland |
| Motor skills domain Standard Score | Vineland |
| Repetitive motor mean frequency score | RBS-R |
| **Domain 2: Social interaction** | |
| Interpersonal relationships subdomain V-scale Score | Vineland |
| Play and leisure time subdomain V-scale Score | Vineland |
| Interpersonal relationships subdomain Raw Score | Vineland |
| Play and leisure time subdomain Raw Score | Vineland |
| Socialization domain V-scale Score | Vineland |
| Socialization domain Standard Score | Vineland |
| Competence domain: compliance (mean raw score) | ITSEA |
| Competence domain: imitation/play (mean raw score) | ITSEA |
| Competence domain: empathy (mean raw score) | ITSEA |
| Internalizing domain: inhibition to novelty (mean raw score) | ITSEA |
| Social relatedness item cluster (mean raw score) | ITSEA |
| Hours spent with other children weekly | ITSEA |
| **Domain 3: Visual / cognitive** | |
| Visual Reception T Score | MSEL |
| Visual Reception Raw Score | MSEL |
| Cognitive T Score Sum | MSEL |
| **Domain 4: Daily living** | |
| Personal subdomain Raw Score | Vineland |
| Domestic subdomain Raw Score | Vineland |
| Community subdomain Raw Score | Vineland |
| Coping skills subdomain Raw Score | Vineland |
| Personal subdomain V-scale Score | Vineland |
| Domestic subdomain V-scale Score | Vineland |
| Community subdomain V-scale Score | Vineland |
| Coping skills subdomain V-scale Score | Vineland |
| Living skills domain V-scale Score | Vineland |
| Living skills domain Standard Score | Vineland |
| Sum of domain standard scores | Vineland |
| **Domain 5: Emotion / regulation** | |
| Total Competence T Score | ITSEA |
| Dysregulation domain: sleep (mean raw score) | ITSEA |
| Dysregulation domain: eating (mean raw score) | ITSEA |
| Competence domain: mastery motivation (mean raw score) | ITSEA |
| Externalizing domain mean raw score | ITSEA |

*Continued on next page*

*Table S2 continued*

| **Measure / score** | **Source instrument** |
|---|---|
| Internalizing domain mean raw score | ITSEA |
| Externalizing domain: activity/impulsivity (mean score) | ITSEA |
| Internalizing domain: depression/withdrawal (mean score) | ITSEA |
| Externalizing domain: aggression/defiance (mean raw score) | ITSEA |
| Externalizing domain: peer aggression (mean raw score) | ITSEA |
| Competence domain: prosocial peer relations (mean raw score) | ITSEA |
| Maladaptive item cluster mean raw score | ITSEA |
| Atypical item cluster mean raw score | ITSEA |
| Self-directed self-injurious mean frequency score | RBS-R |
| Restricted behavior mean frequency score | RBS-R |
| Ritual/routine behavior mean frequency score | RBS-R |
| Composite score mean frequency score | RBS-R |
| **Domain 6: Speech-related outcomes** | |
| Gesture phrases understood | CDI-WG |
| Total vocabulary comprehension | CDI-WG |
| Total vocabulary production | CDI-WG |
| Receptive Language T Score | MSEL |
| Expressive Language T Score | MSEL |
| Receptive Language Raw Score | MSEL |
| Expressive Language Raw Score | MSEL |
| Receptive subdomain Raw Score | Vineland |
| Expressive subdomain Raw Score | Vineland |
| Receptive subdomain V-scale Score | Vineland |
| Expressive subdomain V-scale Score | Vineland |
| Communication domain Standard Score | Vineland |
| Communication domain V-scale Score | Vineland |

CDI-WG, MacArthur–Bates Communicative Development Inventories: Words and Gestures;

MSEL, Mullen Scales of Early Learning;

Vineland, Vineland Adaptive Behavior Scales;

ITSEA, Infant-Toddler Social and Emotional Assessment;

RBS-R, Repetitive Behavior Scale–Revised.

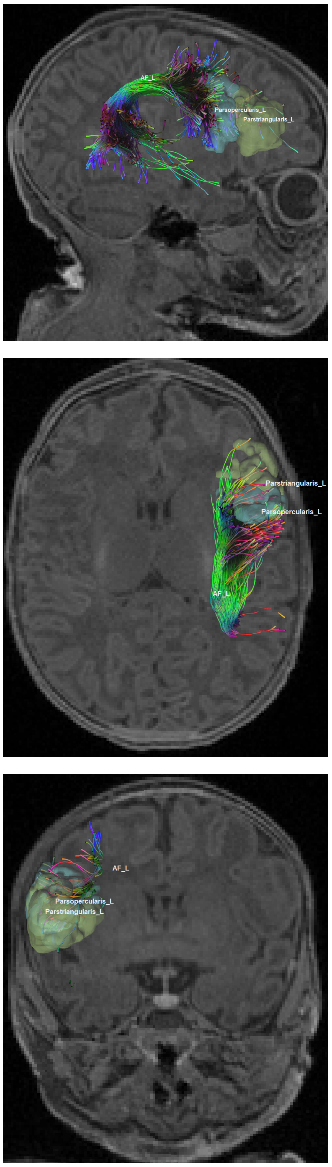


**Figure S4**: **In-brain anatomical views of the neonatal left arcuate fasciculus (AF).** Sagittal, axial, and coronal views are shown in native diffusion space with tract streamlines overlaid on the subject's warped T1w. The neonatal AF terminates predominantly in precentral regions while also showing sparse but plausible extension toward inferior frontal territory, including pars opercularis and pars triangularis. This pattern is broadly consistent with previous reports emphasizing predominant neonatal precentral AF connectivity (*48*, *49*), while illustrating the increased sensitivity of HARDI/CSD-based infant tractography to sparse inferior frontal extensions. This figure is presented as anatomical support for a weak early structural scaffold rather than as evidence of an adult-like language circuit at birth.

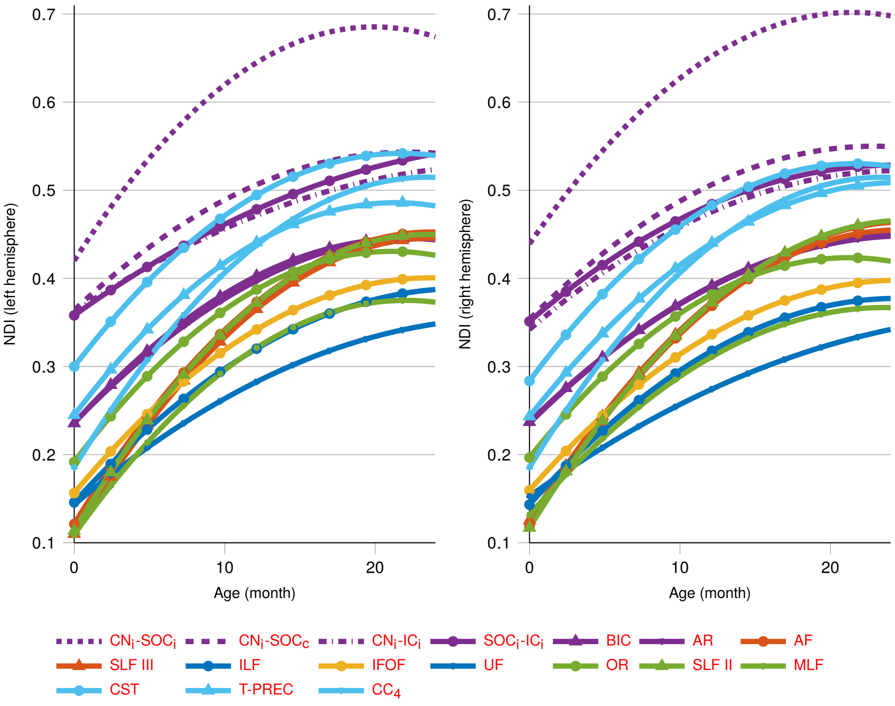


**Figure S5**: **NDI developmental trajectories of tracts across both hemispheres.** Solid lines show fixed-effect predicted NDI trajectories from birth to 24 months for each tract in the left and right hemispheres. Colors denote tract systems as in the main text. This figure complements main-text Figure 3 by extending the trajectory summary to both hemispheres.

**Table S3**: **Model-comparison results for developmental charting of auditory tracts.** Entries compare linear, logarithmic, and quadratic age functions for the trajectory-visualization models. These tables justify the use of quadratic models for developmental curve display in the main text, whereas separate log-age models were used for tract-wise maturation-rate estimation.

| **Tract name** | | $CN_L$-$SOC_L$ | $CN_L$-$SOC_R$ | $CN_R$-$SOC_R$ | $CN_R$-$SOC_L$ | $CN_L$-$IC_L$ | $SOC_L$-$IC_L$ | $CN_R$-$IC_R$ | $SOC_R$-$IC_R$ | $BIC_L$ | $AR_L$ | $BIC_R$ | $AR_R$ |
|---|---|---|---|---|---|---|---|---|---|---|---|---|---|
| **Linear** | $p$ | 4.79e-26 | 1.89e-35 | 1.70e-26 | 3.05e-41 | 1.47e-27 | 1.69e-33 | 1.44e-33 | 6.43e-28 | 3.71e-41 | 6.47e-56 | 2.03e-41 | 1.48e-60 |
| | $r^2 \uparrow$ | 0.7140 | 0.8796 | 0.8074 | 0.8909 | 0.8818 | 0.7923 | 0.7408 | 0.7946 | 0.9068 | 0.8892 | 0.9011 | 0.8476 |
| | AIC ↓ | -514.23 | -816.07 | -519.94 | -836.21 | **-716.33** | **-718.99** | -694.47 | -694.17 | -770.46 | -874.04 | -747.59 | -871.90 |
| **Logarithm** | $p \downarrow$ | 5.04e-36 | 3.79e-46 | 8.38e-39 | 2.30e-53 | 2.21e-27 | 3.49e-31 | 4.74e-31 | 1.37e-36 | 3.23e-43 | 1.83e-56 | 4.35e-39 | 2.93e-55 |
| | $r^2 \uparrow$ | 0.7812 | 0.8794 | 0.8318 | 0.8580 | 0.8395 | 0.7609 | **0.7758** | 0.7900 | 0.9072 | 0.9113 | **0.9141** | 0.8895 |
| | AIC ↓ | **-555.17** | **-847.69** | **-561.43** | **-857.30** | -707.14 | -712.44 | **-703.52** | **-722.32** | **-795.78** | **-906.62** | **-764.61** | **-896.21** |
| **Quadratic** | $p$ | 6.90e-17 | 2.00e-19 | 3.39e-12 | 3.65e-24 | 1.64e-09 | 6.97e-08 | 3.17e-12 | 9.87e-11 | 2.90e-25 | 3.28e-26 | 1.43e-16 | 7.84e-26 |
| | $r^2 \uparrow$ | **0.8211** | **0.9319** | **0.8747** | **0.9032** | **0.9013** | **0.8110** | 0.7675 | **0.8038** | **0.9410** | **0.9254** | 0.9055 | **0.8950** |
| | AIC ↓ | -524.66 | -820.68 | -528.20 | -838.97 | -704.00 | -698.70 | -683.24 | -690.49 | -793.89 | -900.76 | -750.89 | -890.55 |

AIC=akaike information criterion; L=left; R=right; CN=cochlear nucleus; SOC=superior olivary complex; IC=inferior colliculus; BIC=brachium of the inferior colliculus; AR=acoustic radiation.

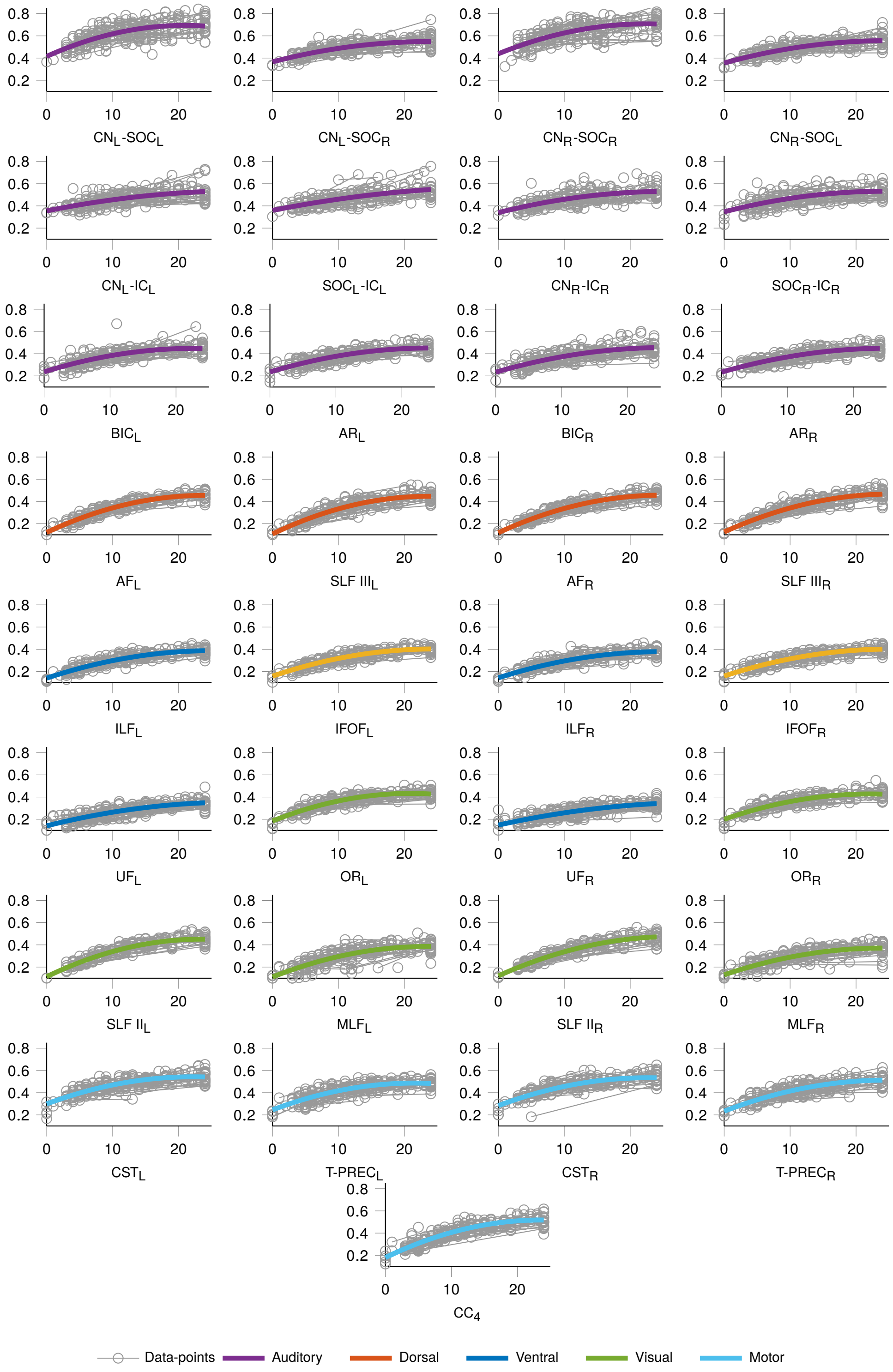


**Figure S6**: **Individual NDI data points and fitted curves across both hemispheres.** Gray circles denote visit-level tract-averaged NDI values, and colored lines denote fixed-effect predicted trajectories from the mixed-effects models. This figure illustrates both inter-individual variability and the overall developmental pattern summarized more compactly in Figure S5.

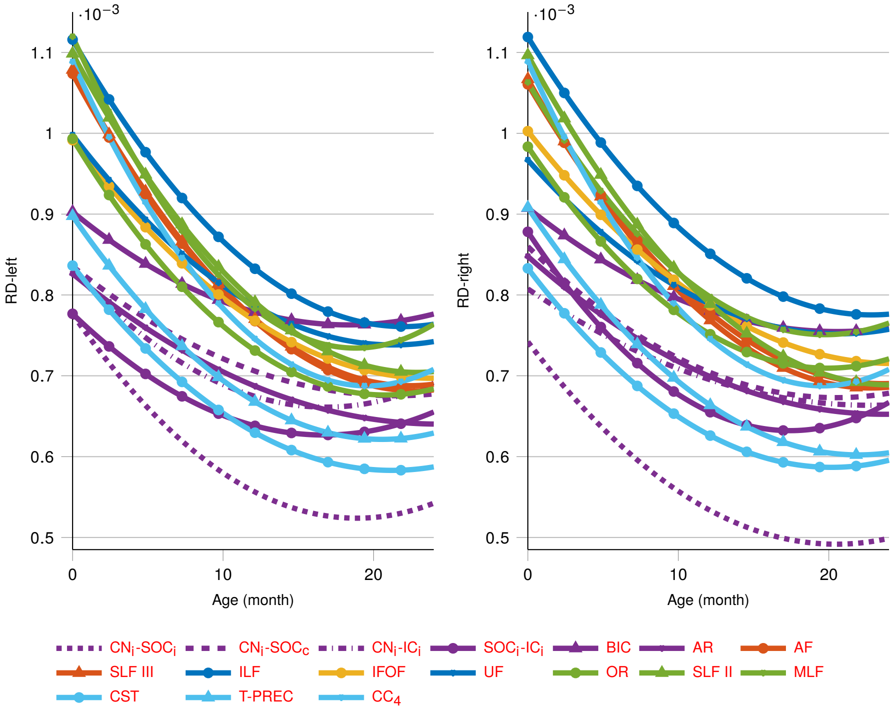


**Figure S7**: **RD developmental trajectories of tracts across both hemispheres.** Solid lines show fixed-effect predicted RD trajectories from birth to 24 months for each tract in the left and right hemispheres. These tensor-derived curves are provided as benchmark analyses for comparison with the primary NODDI-based NDI results.

**Table S4**: **Model-comparison results for developmental charting of major tracts in left hemisphere.** Entries compare linear, logarithmic, and quadratic age functions for the trajectory-visualization models. These tables justify the use of quadratic models for developmental curve display in the main text, whereas separate log-age models were used for tract-wise maturation-rate estimation.

| **Tract name** | | AF | SLF III | ILF | IFOF | UF | OR | SLF II | MLF | CST | T-PREC | $CC_4$ |
|---|---|---|---|---|---|---|---|---|---|---|---|---|
| **Linear** | $p$ | 8.38e-81 | 2.25e-72 | 7.59e-63 | 1.11e-71 | 1.75e-57 | 1.47e-49 | 3.70e-78 | 6.63e-51 | 8.66e-46 | 4.49e-46 | 2.85e-66 |
| | $r^2\uparrow$ | 0.9515 | 0.9426 | 0.9469 | 0.9530 | 0.9549 | 0.9256 | 0.9669 | 0.8200 | 0.9125 | 0.7691 | 0.9193 |
| | AIC $\downarrow$ | -832.38 | -751.84 | -867.01 | -909.85 | -896.60 | -834.53 | -810.49 | -652.56 | -776.55 | -771.07 | -735.43 |
| **Logarithm** | $p$ | 1.90e-90 | 3.62e-77 | 2.67e-67 | 2.21e-75 | 4.83e-48 | 3.56e-76 | 1.42e-100 | 6.21e-43 | 1.34e-68 | 2.58e-55 | 2.21e-67 |
| | $r^2\uparrow$ | 0.9807 | 0.9747 | **0.9561** | **0.9708** | 0.9443 | 0.9565 | 0.9932 | 0.8935 | 0.9236 | 0.8223 | **0.9635** |
| | AIC $\downarrow$ | -936.45 | -818.96 | **-908.26** | -963.81 | -880.99 | **-949.98** | -943.79 | -672.10 | **-850.82** | **-828.91** | -797.66 |
| **Quadratic** | $p$ | 1.10e-71 | 2.80e-53 | 7.62e-42 | 1.48e-47 | 3.18e-20 | 1.12e-52 | 3.65e-71 | 1.36e-22 | 1.32e-27 | 7.00e-32 | 2.05e-45 |
| | $r^2\uparrow$ | **0.9809** | **0.9837** | 0.9233 | 0.9647 | **0.9797** | **0.9628** | **0.9959** | **0.9027** | **0.9379** | **0.8250** | 0.9534 |
| | AIC $\downarrow$ | **-961.34** | **-833.29** | -904.68 | **-979.13** | **-907.79** | -937.37 | **-966.21** | **-673.90** | -819.47 | -816.58 | **-807.99** |

AIC=akaike information criterion; AF: arcuate fasciculus; SLF III: superior longitudinal fasciculus III; ILF: inferior longitudinal fasciculus; IFOF: inferior fronto-occipital fasciculus; UF: uncinate fasciculus; OR: optic radiation; SLF II: superior longitudinal fasciculus II; MLF: middle longitudinal fasciculus; CST: corticospinal tract; T-PREC: thalamo-precentral; and CC 4: corpus callosum anterior midbody (primary motor).

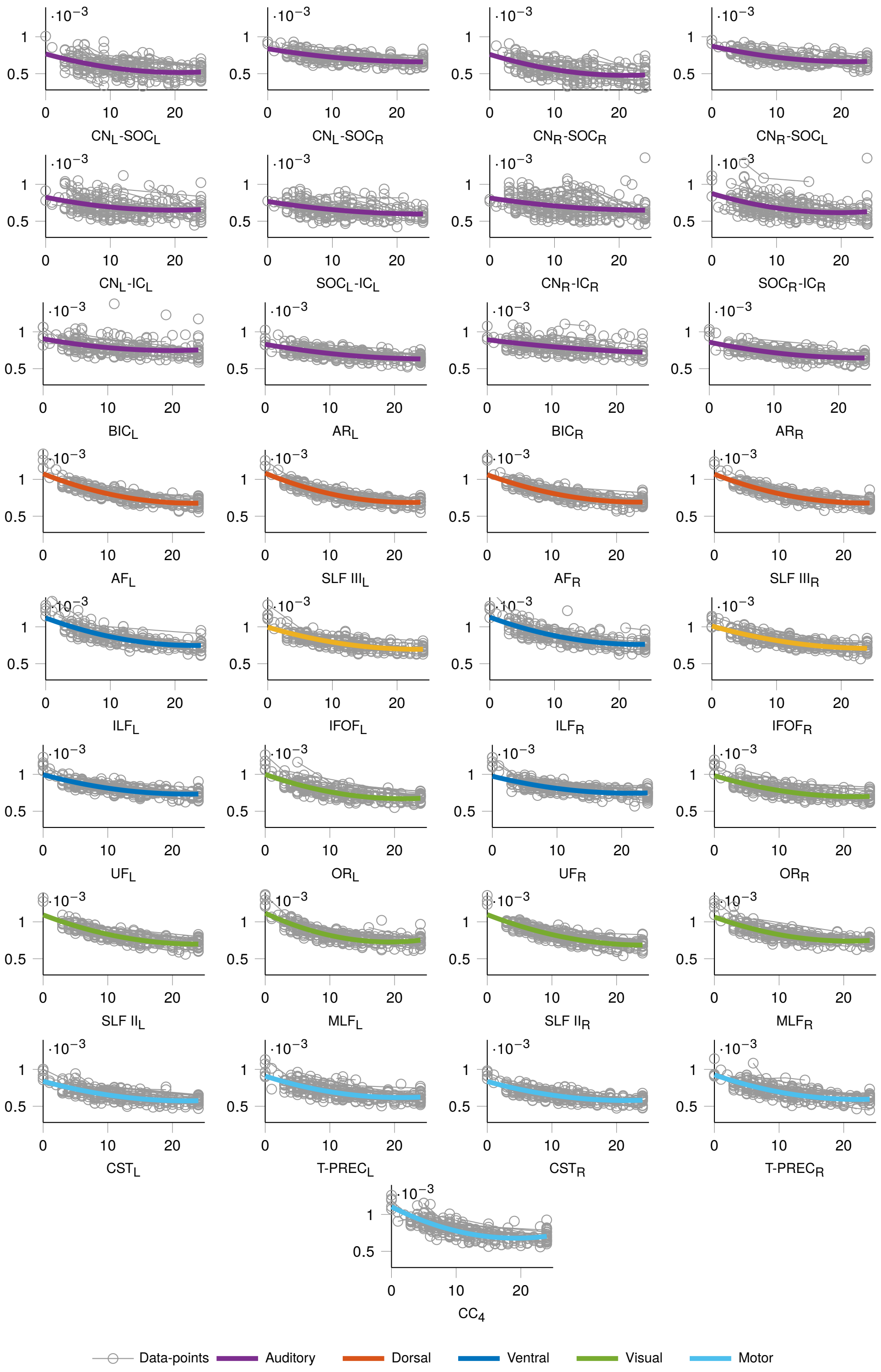


**Figure S8**: **Individual RD data points and fitted curves across both hemispheres.** Gray circles denote visit-level tract-averaged RD values, and colored lines denote fixed-effect predicted trajectories. This figure provides the observation-level counterpart to Figure S7 and is included as a benchmark.

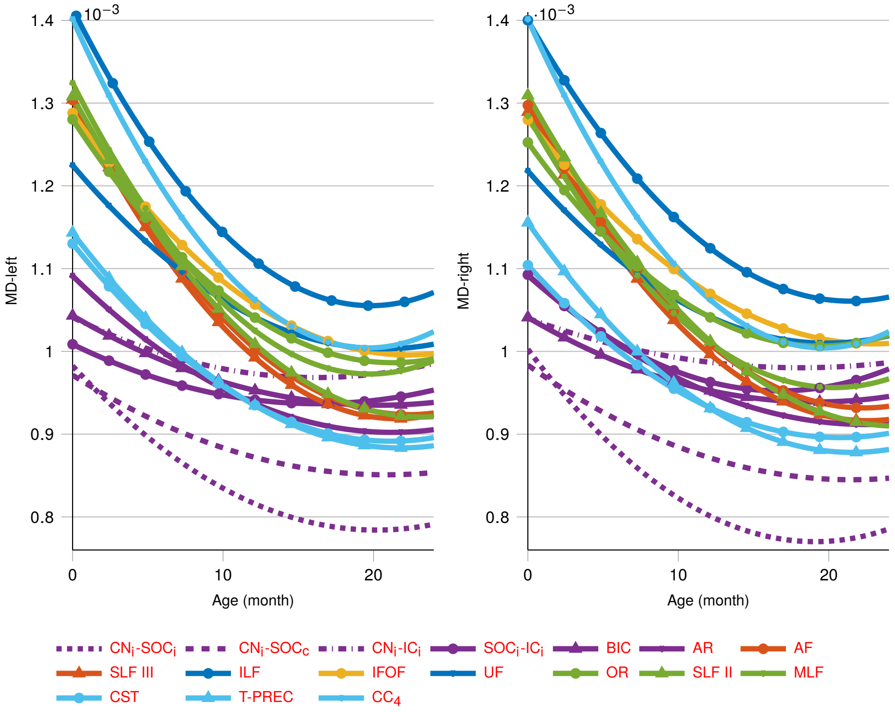


**Figure S9**: **MD developmental trajectories of tracts across both hemispheres.** Solid lines show fixed-effect predicted MD trajectories from birth to 24 months for each tract in the left and right hemispheres. These tensor-derived results are shown for benchmarking relative to the primary NDI findings.

**Table S5**: **Model-comparison results for developmental charting of tracts in right-hemisphere .** Entries compare linear, logarithmic, and quadratic age functions for the trajectory-visualization models. These comparisons were used to justify the quadratic models employed for developmental curve display in the main text, whereas separate log-age models were used for tract-wise maturation-rate estimation.

| Tract name | | AF | SLF III | ILF | IFOF | UF | OR | SLF II | MLF | CST | T-PREC | $CC_4$ |
|---|---|---|---|---|---|---|---|---|---|---|---|---|
| **Linear** | $p$ | 1.15e-86 | 1.87e-85 | 2.48e-63 | 1.17e-68 | 6.98e-53 | 3.54e-50 | 1.40e-88 | 1.26e-49 | 1.15e-86 | 1.70e-62 | 2.85e-66 |
| | $r^2$ ↑ | 0.9494 | 0.9386 | 0.9208 | 0.9354 | 0.9410 | 0.9688 | 0.9766 | 0.9211 | 0.9279 | 0.8413 | 0.9193 |
| | AIC ↓ | -873.23 | -827.82 | -843.14 | -918.38 | -912.78 | -839.25 | -865.23 | -793.70 | -776.44 | -774.12 | -735.43 |
| **Logarithm** | $p$ | 3.03e-94 | 9.26e-85 | 9.14e-63 | 3.21e-70 | 2.48e-44 | 1.56e-77 | 3.32e-93 | 2.53e-51 | 3.03e-94 | 1.78e-59 | 2.21e-67 |
| | $r^2$ ↑ | 0.9769 | **0.9694** | **0.9348** | 0.9613 | 0.9127 | 0.9584 | 0.9848 | 0.9229 | 0.9197 | 0.9256 | **0.9635** |
| | AIC ↓ | -963.30 | -882.94 | -869.48 | -962.88 | -888.58 | **-923.97** | -924.86 | **-820.15** | **-837.81** | -811.35 | -797.66 |
| **Quadratic** | $p$ | 2.27e-66 | 4.00e-52 | 3.82e-40 | 1.94e-40 | 4.15e-15 | 5.15e-37 | 8.98e-60 | 4.04e-31 | 2.27e-66 | 3.70e-27 | 2.05e-45 |
| | $r^2$ ↑ | **0.9774** | 0.9686 | 0.9207 | **0.9726** | **0.9812** | **0.9751** | **0.9878** | **0.9580** | **0.9543** | **0.9462** | 0.9534 |
| | AIC ↓ | **-981.34** | **-899.23** | **-877.80** | **-968.36** | **-916.90** | -897.93 | **-946.34** | -818.08 | -822.88 | **-816.36** | **-807.99** |

AIC=akaike information criterion; AF: arcuate fasciculus; SLF III: superior longitudinal fasciculus III; ILF: inferior longitudinal fasciculus; IFOF: inferior fronto-occipital fasciculus; UF: uncinate fasciculus; OR: optic radiation; SLF II: superior longitudinal fasciculus II; MLF: middle longitudinal fasciculus; CST: corticospinal tract; T-PREC: thalamo-precentral; and $CC_4$: corpus callosum anterior midbody (primary motor).

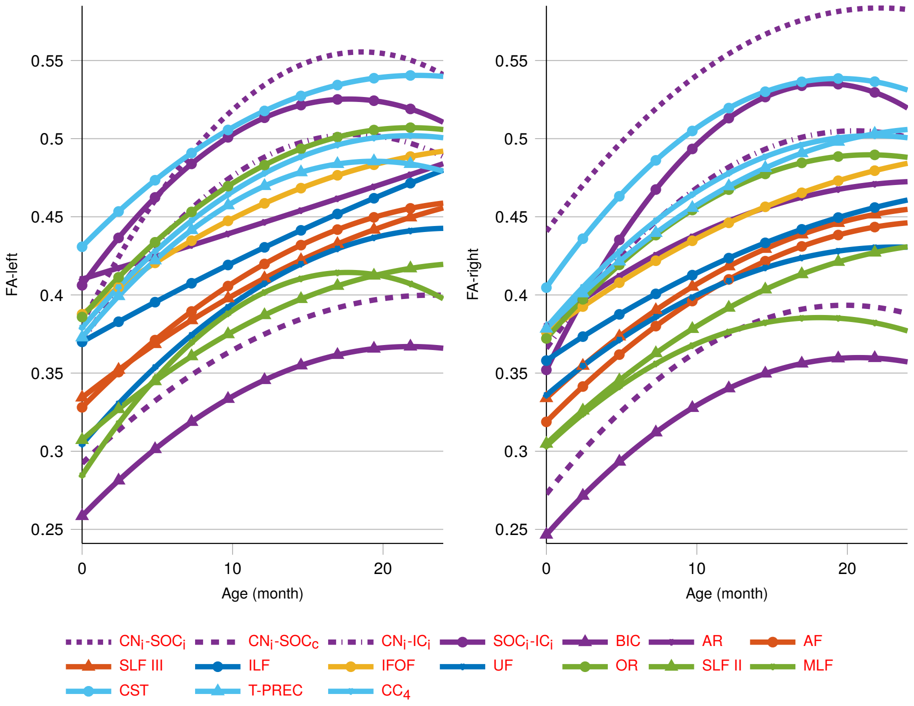


**Figure S10**: **FA developmental trajectories of tracts across both hemispheres.** Solid lines show fixed-effect predicted FA trajectories from birth to 24 months for each tract in the left and right hemispheres. These tensor-derived results are displayed as supplementary benchmarks and are not the primary basis for inference.

**Table S6**: **Compact summary of gradient metrics significantly associated with association tracts.** For each gradient predictor, the table lists the association tracts that survived BH–FDR correction within the predictor family. This compact table corresponds to the tract-level counts summarized in Figure S13.

| Predictor | Significant association-tract outcomes |
|---|---|
| $\nabla_{R}^{M-AR}$ | SLF II$_L$, SLF III$_L$, AF$_L$, ILF$_L$, IFOF$_L$, UF$_L$, MLF$_L$, AF$_R$, SLF III$_R$, SLF II$_R$, ILF$_R$, IFOF$_R$, UF$_R$, MLF$_R$ |
| $\nabla_{L}^{M-AR}$ | AF$_L$, SLF III$_L$, SLF II$_L$, ILF$_L$, IFOF$_L$, UF$_L$, MLF$_L$, AF$_R$, SLF III$_R$, SLF II$_R$, ILF$_R$, IFOF$_R$, UF$_R$ |
| $\nabla_{R}^{M-OR}$ | AF$_L$, SLF III$_L$, SLF II$_L$, ILF$_L$, IFOF$_L$, UF$_L$, MLF$_L$, AF$_R$, SLF III$_R$, SLF II$_R$, UF$_R$ |
| $\nabla_{L}^{M-OR}$ | SLF III$_L$, SLF II$_L$, UF$_L$, MLF$_L$, AF$_R$, SLF III$_R$, SLF II$_R$, UF$_R$ |
| $\nabla_{L}^{AR-OR}$ | ILF$_L$, ILF$_R$ |
| $\nabla_{R}^{AR-OR}$ | None |

Predictors denote the sensory–motor gradient metrics defined in the Methods. This compact table summarizes the tract-level associations counted in Supplementary Figure S13 **b**.

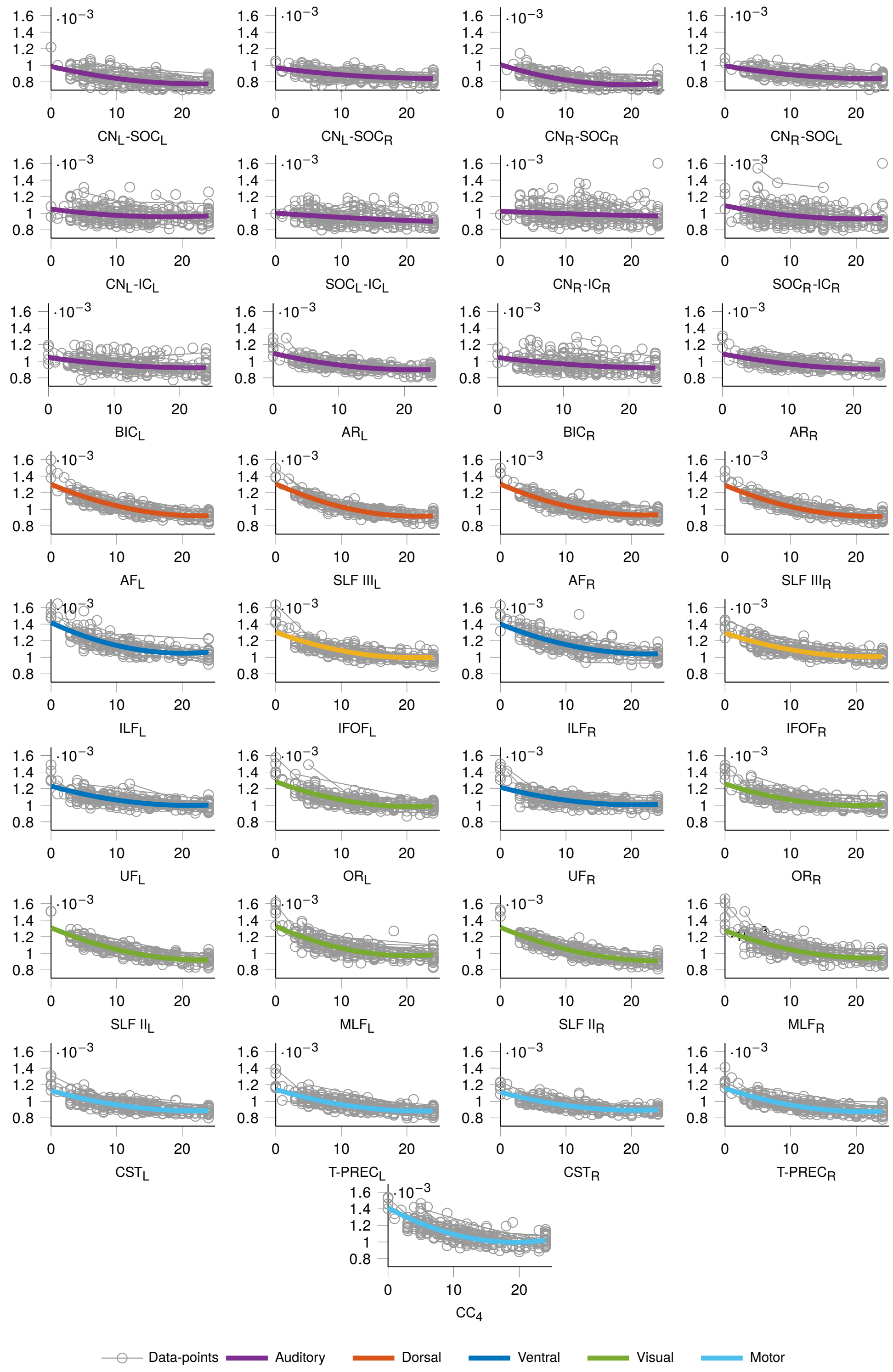


**Figure S11**: **Individual MD data points and fitted curves across both hemispheres.** Gray circles denote visit-level tract-averaged MD values, and colored lines denote fixed-effect predicted trajectories. This figure provides the observation-level counterpart to Figure S9.

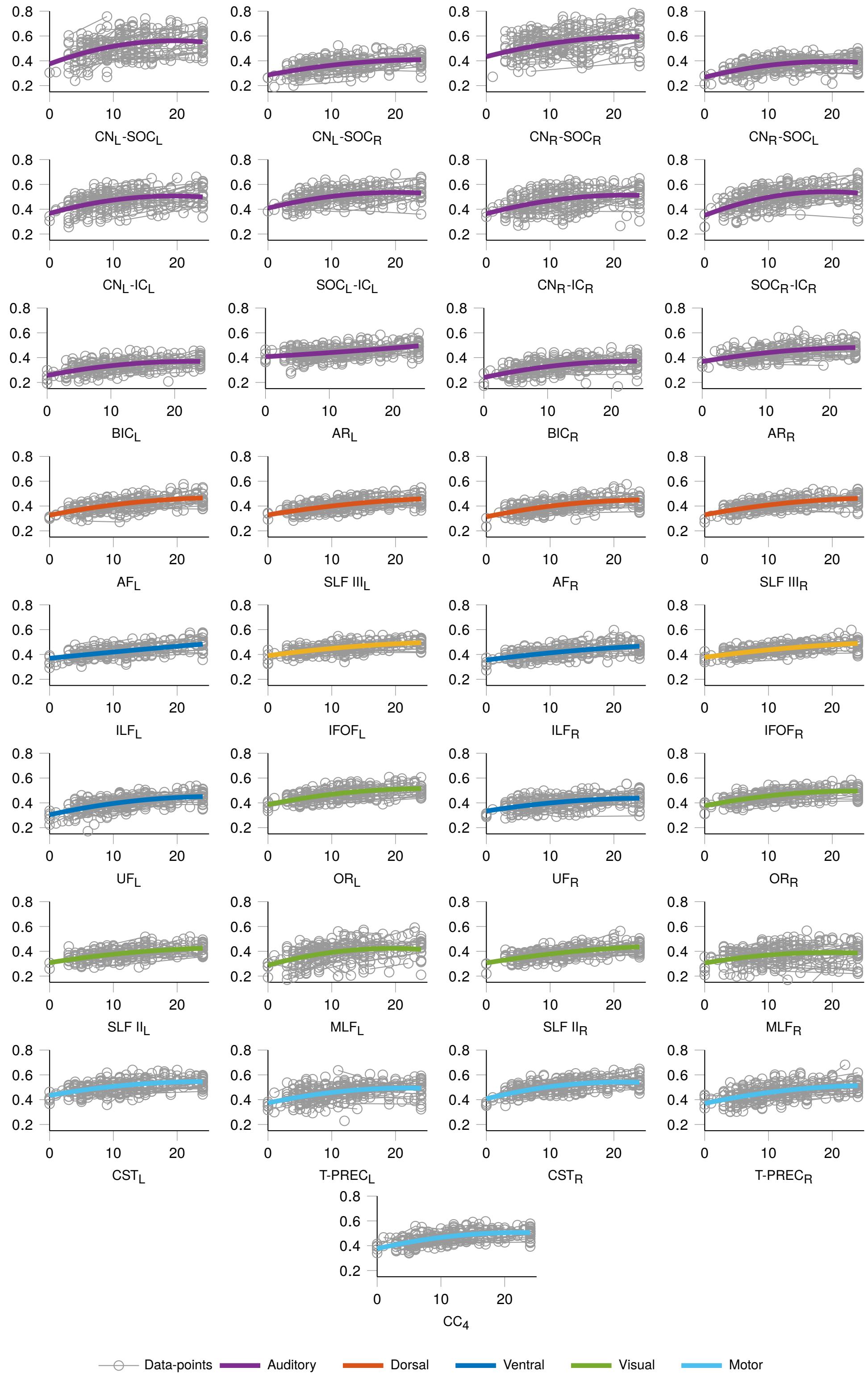


**Figure S12**: **Individual FA data points and fitted curves across both hemispheres.** Gray circles denote visit-level tract-averaged FA values, and colored lines denote fixed-effect predicted trajectories. This figure provides the observation-level counterpart to Figure S10 and illustrates the greater variability of tensor-derived benchmark metrics relative to the primary NDI results.

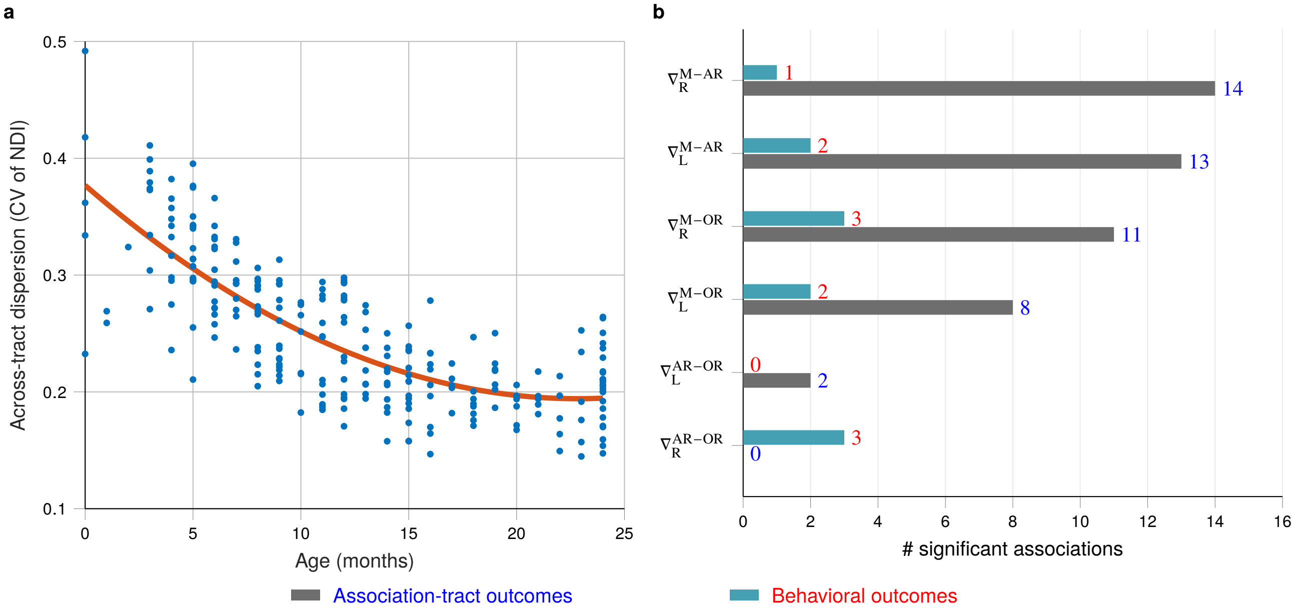


**Figure S13**: **System-level balance analyses supporting progressive cross-system rebalancing.** (**a**) Inter-tract dispersion, quantified as the coefficient of variation (CV) of tract-level NDI across the representative tract set, decreases with age. Each point denotes one scan, and the solid curve shows the fitted trajectory from the quadratic mixed-effects model described in Eqn. S2. Lower CV indicates a less heterogeneous tract-maturity profile across systems. (**b**) Summary counts of significant associations for sensory–motor gradient metrics. Rows show visit-wise gradients defined relative to a motor anchor ($CC_4$) and sensory anchors (AR, OR), separately by hemisphere; bars indicate the numbers of significant associations with association-tract and behavioral outcomes after BH–FDR correction within each predictor family. Motor-leading gradients ($CC_4$ relative to AR/OR) showed substantially more tract-level associations than sensory–sensory contrasts between OR and AR.

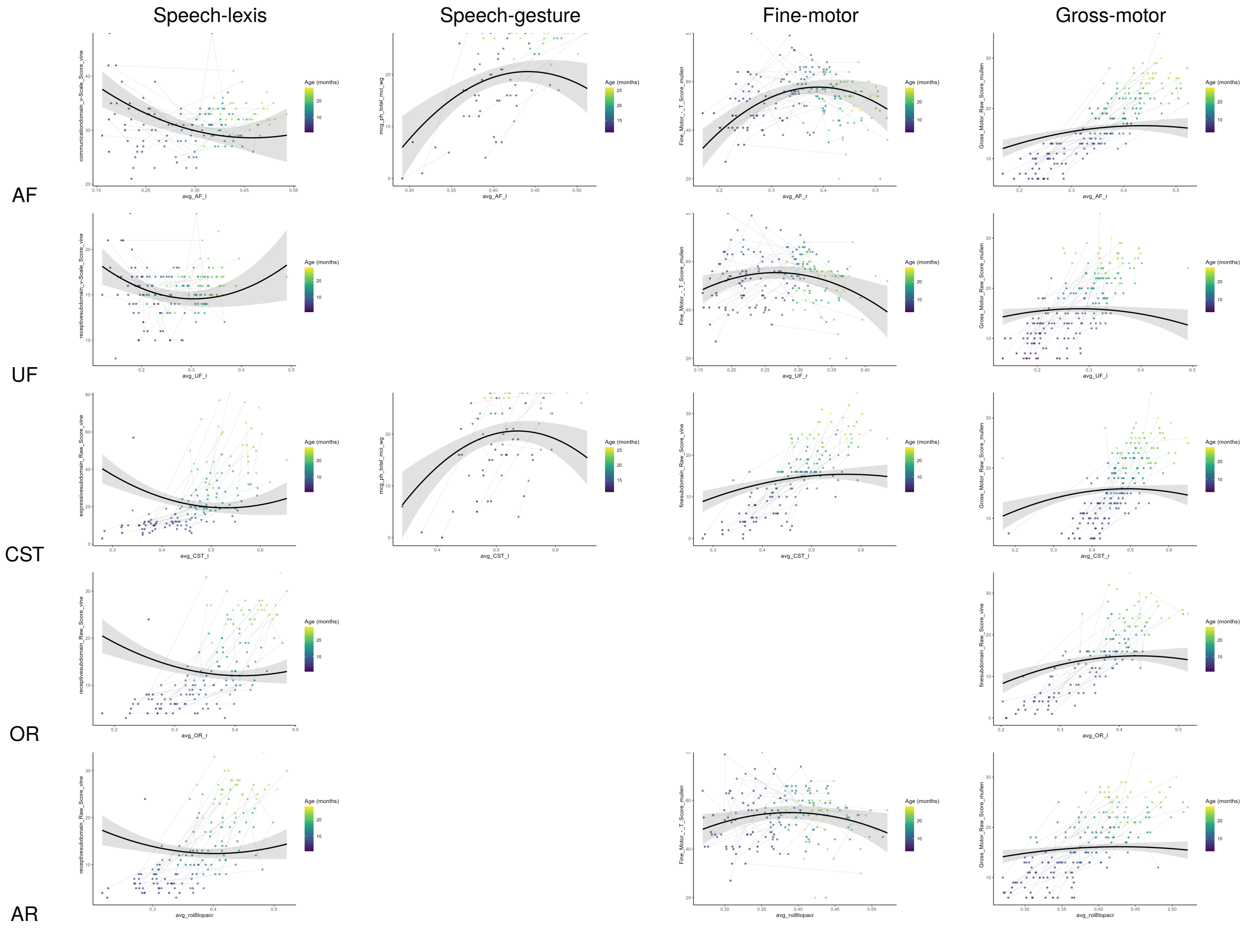


**Figure S14**: **Representative tract–behavior associations illustrating positive, negative, and curvilinear patterns.** Each panel shows one tract–behavior pair that survived BH–FDR correction ($q < 0.05$) in the screening analysis. Each point denotes one visit and is colored by age (months); thin gray lines connect repeated visits within infant. Black curves and shaded bands show the age- and sex-adjusted population-level marginal association estimated from the mixed-effects model, $Outcome \sim 1 + NDI + NDI2 + Age + Sex + (1 \mid Subject)$, with subject-specific random effects set to zero and shaded bands denoting 95% confidence intervals. Panels were selected to exemplify the principal patterns summarized in main-text Figure 4

**Table S7**: **Behavioral outcomes significantly associated with sensory–motor gradient metrics.** For each retained association, the table reports the gradient predictor, behavioral outcome, mixed-effects coefficient ($\beta$), FDR-adjusted $q$ value, and sample size ($N_{\text{obs}}$: visits; $N_{\text{sub}}$: infants). Models included age, sex, and repeated visits as covariates. Associations from low-$N$ subsets, particularly CBCL- and ITSEA-derived outcomes, are interpreted as exploratory even when meeting the FDR threshold.

| **Predictor** | **Outcome** | ***Estimate*** | $q$ | $N_{\text{obs}}$ | $N_{\text{sub}}$ |
|---|---|---|---|---|---|
| $\nabla_{\text{L}}^{\text{M-AR}}$ | Fine Motor raw score (MSEL) | 20.91 | 0.009 | 179 | 121 |
| $\nabla_{\text{L}}^{\text{M-AR}}$ | Gross Motor raw score (MSEL) | 16.76 | 0.036 | 179 | 121 |
| $\nabla_{\text{R}}^{\text{M-AR}}$ | Fine Motor raw score (MSEL) | 21.16 | 0.014 | 177 | 119 |
| $\nabla_{\text{L}}^{\text{M-OR}}$ | Gross subdomain raw score (Vineland) | 70.29 | 0.019 | 137 | 93 |
| $\nabla_{\text{L}}^{\text{M-OR}}$ | Fine Motor raw score (MSEL) | 24.06 | 0.048 | 168 | 119 |
| $\nabla_{\text{R}}^{\text{M-OR}}$ | Fine Motor raw score (MSEL) | 23.82 | 0.042 | 163 | 116 |
| $\nabla_{\text{R}}^{\text{M-OR}}$ | Affective Problems raw score (CBCL) | -63.47 | 0.042 | 39 | 37 |
| $\nabla_{\text{R}}^{\text{M-OR}}$ | Affective Problems T score (CBCL) | -125.6 | 0.042 | 39 | 37 |
| $\nabla_{\text{R}}^{\text{AR-OR}}$ | Attention Problems raw score (CBCL) | -15.55 | $< 0.001$ | 39 | 37 |
| $\nabla_{\text{R}}^{\text{AR-OR}}$ | Attention Deficit/Hyperactivity raw (CBCL) | -15.23 | 0.022 | 39 | 37 |
| $\nabla_{\text{R}}^{\text{AR-OR}}$ | Total Competence T score (ITSEA) | 101.89 | 0.031 | 79 | 64 |

$\nabla_{\text{L}}^{\text{M-AR}}$, $\nabla_{\text{R}}^{\text{M-AR}}$, $\nabla_{\text{L}}^{\text{M-OR}}$, $\nabla_{\text{R}}^{\text{M-OR}}$, $\nabla_{\text{L}}^{\text{AR-OR}}$, and $\nabla_{\text{R}}^{\text{AR-OR}}$denote the gradient metrics defined in the Eqn. S3.

Associations from low-$N$ subsets, particularly CBCL- and ITSEA-derived outcomes, are interpreted as exploratory even when meeting the FDR threshold.

**Table S8**: **Summary of significant cross-sectional mediation pathway families identified via piecewise structural equation modeling.** Candidate developmental pathways were screened across $74{,}409$ specified triplets using tract-specific mean NDI values and behavioral measures. Of these, $74{,}276$ triplets were testable and $849$ were retained after study-wide BH–FDR correction. For each pathway family, the table reports the triplet structure, the number of testable triplets, the number retained after correction, and the corresponding developmental rationale. These results are presented as a prioritization and hypothesis-generation step rather than as evidence of causal mediation.

| ID | Pathway model | Triplet | $N_{\text{sig}}$ | Purpose and theoretical hypothesis |
|---|---|---|---|---|
| A1 | *Tract → Motor → Speech* | 6,006 | 44 | Tract supports speech indirectly via enhanced motor skills. |
| A2 | *Tract → Social → Speech* | 5,148 | 0 | Tract supports speech indirectly via enhanced social interactions. |
| A3 | *Tract → Visual → Speech* | 1,287 | 7 | Tract supports speech indirectly via visual processing. |
| B1 | *Tract → Social → Motor* | 5,544 | 0 | Tract supports motor skills indirectly via enhanced social interactions. |
| B2 | *Tract → Motor → Social* | 5,544 | 42 | Tract supports social interactions indirectly via enhanced motor skills. |
| B3 | *Tract → Motor → Emotional* | 7,854 | 0 | Tract supports emotional regulation indirectly via motor skills. |
| B4 | *Tract → Social → Emotional* | 6,732 | 0 | Tract supports emotional regulation indirectly via social interactions. |
| B5 | *Tract → Motor → Living Skills* | 5,082 | 24 | Tract supports living skills indirectly via enhanced motor skills. |
| B6 | *Tract → Social → Living Skills* | 4,356 | 0 | Tract supports living skills indirectly via enhanced social interactions. |
| C1 | *Tract → Emotional → Speech* | 7,293 | 0 | Tract supports speech indirectly via enhanced emotional regulation. |
| C2 | *Tract → Living Skills → Speech* | 4,719 | 1 | Tract supports speech indirectly via enhanced living skills. |
| D1 | *Motor → Visual → Speech* | 546 | 80 | Motor skills support speech indirectly via improved visual processing. |
| D2 | *Social → Visual → Speech* | 468 | 3 | Social interactions support speech indirectly via improved visual processing. |
| D3 | *Motor → Emotional → Speech* | 3,094 | 4 | Motor skills support speech indirectly via improved emotional regulation. |
| D4 | *Social → Emotional → Speech* | 2,652 | 1 | Social interactions support speech indirectly via improved emotional regulation. |
| D5 | *Motor → Living Skills → Speech* | 2,002 | 171 | Motor skills support speech indirectly via improved living skills. |
| D6 | *Social → Living Skills → Speech* | 1,716 | 164 | Social interactions support speech indirectly via improved living skills. |
| E1 | *Social → Motor → Speech* | 2,184 | 145 | Social interactions facilitate speech indirectly through enhanced motor skills. |
| E2 | *Motor → Social → Speech* | 2,184 | 163 | Motor skills facilitate speech indirectly through enhanced social interactions. |
| | $74,409$ **Combinations in Total** | | | $849$ **out of** $74,276$ **potential pathway using FDR correction** |

**Table S9**: **Summary of significant longitudinal mediation pathways from motor skills to speech-related outcomes via social interaction (*Motor* → *Social* → *Speech*).** Predictor variables (motor skills) were assessed at approximately 6 months, mediator variables (social-interaction measures) at approximately 12 months, and outcome variables (speech-related measures) at approximately 24 months using matched longitudinal data from the BCP cohort. Entries report path estimates, standard errors, and uncorrected $p$ values for the *Motor* → *Social* and *Social* → *Speech* components estimated within the piecewise structural equation modeling framework. Because these analyses were conducted in modest complete-data subsets without FDR correction, the results are interpreted as exploratory and hypothesis-generating rather than causal.

| **Predictor (Motor)** | **Mediator (Social)** | **Outcome (Speech)** | **Motor → Social** | | | **Social → Speech** | | |
|---|---|---|---|---|---|---|---|---|
| | | | *Estimate* | *Std. Error* | $p$-value | *Estimate* | *Std. Error* | $p$-value |
| grosssubdomain Raw | interpersrelationsubdom Raw | mcg vc totcom | 0.711 | 0.238 | $0.003^{**}$ | 8.535 | 4.343 | $0.049^{*}$ |
| grosssubdomain Raw | interpersrelationsubdom Raw | mcg vc totpr | 0.711 | 0.238 | $0.003^{**}$ | 18.566 | 7.190 | $0.010^{*}$ |
| grosssubdomain Raw | interpersrelationsubdom Raw | Receptive Language Raw | 0.644 | 0.227 | $0.005^{**}$ | 0.562 | 0.230 | $0.014^{*}$ |
| grosssubdomain Raw | interpersrelationsubdom Raw | Receptive Language T | 0.644 | 0.227 | $0.005^{**}$ | 1.552 | 0.672 | $0.021^{*}$ |
| grosssubdomain Raw | interpersrelationsubdom Raw | receptive subdomain Raw | 0.620 | 0.219 | $0.005^{**}$ | 0.475 | 0.235 | $0.043^{*}$ |
| grosssubdomain Raw | interpersrelationsubdom Raw | receptive subdomain v Scale | 0.620 | 0.219 | $0.005^{**}$ | 0.289 | 0.146 | $0.048^{*}$ |
| motor skills domain Standard | interpersrelation subdom Raw | mcg vc totpr | 0.121 | 0.036 | $0.001^{***}$ | 16.542 | 7.516 | $0.028^{*}$ |
| motor skills domain v Scale | interpersrelation subdom Raw | mcg vc totpr | 0.443 | 0.125 | $0.0004^{***}$ | 16.653 | 7.603 | $0.028^{*}$ |
| grosssubdomain Raw | interpersrelation subdom v Scale | mcg vc totcom | 0.574 | 0.176 | $0.001^{**}$ | 11.610 | 5.923 | $0.050^{*}$ |
| grosssubdomain Raw | interpersrelation subdom v Scale | mcg vc totpr | 0.574 | 0.176 | $0.001^{**}$ | 23.284 | 9.956 | $0.019^{*}$ |
| grosssubdomain Raw | interpersrelation subdom v Scale | Receptive Language Raw | 0.493 | 0.172 | $0.004^{**}$ | 0.634 | 0.312 | $0.043^{*}$ |
| grosssubdomain v Scale | interpersrelation subdom v Scale | mcg vc totcom | 0.397 | 0.195 | $0.042^{*}$ | 11.704 | 5.439 | $0.031^{*}$ |
| grosssubdomain v Scale | interpersrelation subdom v Scale | mcg vc totpr | 0.397 | 0.195 | $0.042^{*}$ | 20.824 | 9.133 | $0.023^{*}$ |
| grosssubdomain Raw | socialization domain Standard | mcg vc totpr | 2.110 | 0.781 | $0.007^{**}$ | 5.741 | 2.277 | $0.012^{*}$ |
| motorskillsdomain Standard | socialization domain Standard | mcg vc totpr | 0.338 | 0.120 | $0.005^{**}$ | 4.877 | 2.322 | $0.036^{*}$ |
| motorskillsdomain v Scale | socialization domain Standard | mcg vc totpr | 1.269 | 0.418 | $0.002^{**}$ | 4.912 | 2.349 | $0.037^{*}$ |

Note: Significance levels indicated by: $^{*}p < 0.05$, $^{**}p < 0.01$, $^{***}p < 0.001$.

**Table S10**: **Summary of significant longitudinal mediation pathways from social interaction to speech-related outcomes via motor skills (*Social* → *Motor* → *Speech*).** Predictor variables (social-interaction measures) were assessed at approximately 6 months, mediator variables (motor skills) at approximately 12 months, and outcome variables (speech-related measures) at approximately 24 months using matched longitudinal data from the BCP cohort. Entries report path estimates, standard errors, and uncorrected $p$ values for the *Social* → *Motor* and *Motor* → *Speech* components estimated within the piecewise structural equation modeling framework. Because these analyses were conducted in modest complete-data subsets without FDR correction, the results are interpreted as exploratory and hypothesis-generating rather than causal.

| **Predictor (Social)** | **Mediator (Motor)** | **Outcome (Speech)** | **Social → Motor** | | | **Motor → Speech** | | |
|---|---|---|---|---|---|---|---|---|
| | | | *Estimate* | *Std. Error* | $p$-value | *Estimate* | *Std. Error* | $p$-value |
| interpersrelationsubdom Raw | mcg lg total | Expressive Language Raw | 0.790 | 0.355 | $0.026^{*}$ | 0.401 | 0.146 | $0.006^{**}$ |
| interpersrelationsubdom Raw | mcg lg total | Expressive Language T | 0.790 | 0.355 | $0.026^{*}$ | 1.018 | 0.370 | $0.006^{**}$ |
| interpersrelationsubdom Raw | mcg lg total | mcg vc totpr | 0.927 | 0.331 | $0.005^{**}$ | 7.597 | 3.676 | $0.039^{*}$ |
| interpersrelationsubdom Raw | mcg tg total | Expressive Language Raw | 0.923 | 0.448 | $0.039^{*}$ | 0.318 | 0.114 | $0.005^{**}$ |
| interpersrelationsubdom Raw | mcg tg total | Expressive Language T | 0.923 | 0.448 | $0.039^{*}$ | 0.808 | 0.287 | $0.005^{**}$ |
| interpersrelationsubdom Raw | motor skills domain Standard | mcg ph total | 1.255 | 0.536 | $0.019^{*}$ | 0.073 | 0.035 | $0.038^{*}$ |
| interpersrelationsubdom v Scale | mcg lg total | Expressive Language Raw | 1.199 | 0.583 | $0.040^{*}$ | 0.375 | 0.147 | $0.011^{*}$ |
| interpersrelationsubdom v Scale | mcg lg total | Expressive Language T | 1.199 | 0.583 | $0.040^{*}$ | 0.959 | 0.371 | $0.010^{*}$ |
| interpersrelationsubdom v Scale | mcg lg total | mcg vc totpr | 1.356 | 0.522 | $0.009^{**}$ | 7.209 | 3.602 | $0.045^{*}$ |
| interpersrelationsubdom v Scale | motor skills domain Standard | mcg ph total | 1.808 | 0.902 | $0.045^{*}$ | 0.077 | 0.035 | $0.025^{*}$ |
| playleisuretimesubdom Raw | finesubdomain v Scale | Receptive Language Raw | 0.483 | 0.156 | $0.002^{**}$ | 0.646 | 0.309 | $0.037^{*}$ |
| playleisuretimesubdom Raw | fine subdomain v Scale | Receptive Language T | 0.483 | 0.156 | $0.002^{**}$ | 1.799 | 0.903 | $0.046^{*}$ |
| playleisuretimesubdom Raw | motor skills domain Standard | Receptive Language Raw | 1.925 | 0.961 | $0.045^{*}$ | 0.121 | 0.050 | $0.015^{*}$ |
| playleisuretimesubdom Raw | motor skills domain Standard | Receptive Language T | 1.925 | 0.961 | $0.045^{*}$ | 0.337 | 0.145 | $0.020^{*}$ |
| playleisuretimesubdom v Scale | Fine Motor T | Expressive Language Raw | 1.043 | 0.485 | $0.031^{*}$ | −0.263 | 0.130 | $0.043^{*}$ |
| playleisuretimesubdom v Scale | Fine Motor T | Expressive Language T | 1.043 | 0.485 | $0.031^{*}$ | −0.709 | 0.327 | $0.030^{*}$ |
| playleisuretimesubdom v Scale | finesubdomain v Scale | Receptive Language Raw | 0.379 | 0.124 | $0.002^{**}$ | 0.620 | 0.308 | $0.044^{*}$ |
| playleisuretimesubdom v Scale | motorskills domain Standard | Receptive Language Raw | 1.564 | 0.764 | $0.041^{*}$ | 0.118 | 0.050 | $0.017^{*}$ |
| playleisuretimesubdom v Scale | motorskills domain Standard | Receptive Language T | 1.564 | 0.764 | $0.041^{*}$ | 0.328 | 0.145 | $0.024^{*}$ |
| socializationdomain Standard | Fine Motor T | Expressive Language T | 0.282 | 0.127 | $0.026^{*}$ | −0.710 | 0.332 | $0.032^{*}$ |
| socializationdomain Standard | motorskills domain Standard | Receptive Language Raw | 0.359 | 0.182 | $0.048^{*}$ | 0.113 | 0.049 | $0.022^{*}$ |
| socialization domain Standard | motorskills domain Standard | Receptive Language T | 0.359 | 0.182 | $0.048^{*}$ | 0.311 | 0.144 | $0.032^{*}$ |
| socializationdomain v Scale | Fine Motor T | Expressive Language T | 0.832 | 0.373 | $0.026^{*}$ | −0.707 | 0.332 | $0.033^{*}$ |
| socializationdomain v Scale | motor skills domain Standard | Receptive Language Raw | 1.103 | 0.538 | $0.040^{*}$ | 0.114 | 0.049 | $0.022^{*}$ |
| socializationdomain v Scale | motor skills domain Standard | Receptive Language T | 1.103 | 0.538 | $0.040^{*}$ | 0.312 | 0.145 | $0.031^{*}$ |

Note: Significance levels indicated by: $^{*}p < 0.05$, $^{**}p < 0.01$.

**Table S11**: **Regions of interest (ROIs) defined along the auditory pathway.** The table maps ROI labels to their corresponding anatomical regions along the peripheral, brainstem, thalamic, and cortical auditory pathway. These ROI definitions were used to construct auditory tract variables in Table S12 and in the `Supplementary Data`.

| ROI | Brain region |
|---|---|
| roi1 | cochlear nucleus (CN) left |
| roi2 | cochlear nucleus right |
| roi3 | superior olivary complex (SOC) left |
| roi4 | superior olivary complex right |
| roi5 | inferior colliculus (IC) left |
| roi6 | inferior colliculus right |
| roi7 | medial geniculate body (MGB) left |
| roi8 | medial geniculate body right |
| pacl | primary auditory cortex (PAC) left |
| pacr | primary auditory cortex right |

**Table S12**: **Tract variables used in `Supplementary Data` and downstream analyses.** Entries map each variable name to its source as tract-extracted mean NDI. ROI-to-ROI auditory variables correspond to the auditory-pathway ROIs defined in Table S11 and provide a compact variable dictionary for the `Supplementary Data` and merged analysis tables.

| Variable | Source |
|---|---|
| avg_AF_l | Tract Extraction (NDI mean) |
| avg_AF_r | Tract Extraction (NDI mean) |
| avg_SLF3_l | Tract Extraction (NDI mean) |
| avg_SLF3_r | Tract Extraction (NDI mean) |
| avg_ILF_l | Tract Extraction (NDI mean) |
| avg_ILF_r | Tract Extraction (NDI mean) |
| avg_IFO_l | Tract Extraction (NDI mean) |
| avg_IFO_r | Tract Extraction (NDI mean) |
| avg_UF_l | Tract Extraction (NDI mean) |
| avg_UF_r | Tract Extraction (NDI mean) |
| avg_OR_l | Tract Extraction (NDI mean) |
| avg_OR_r | Tract Extraction (NDI mean) |
| avg_SLF2_l | Tract Extraction (NDI mean) |
| avg_SLF2_r | Tract Extraction (NDI mean) |
| avg_MLF_l | Tract Extraction (NDI mean) |
| avg_MLF_r | Tract Extraction (NDI mean) |
| avg_CST_l | Tract Extraction (NDI mean) |
| avg_CST_r | Tract Extraction (NDI mean) |
| avg_TPREC_l | Tract Extraction (NDI mean) |
| avg_TPREC_r | Tract Extraction (NDI mean) |
| avg_CC_4 | Tract Extraction (NDI mean) |
| avg_roi1toroi4 | Tract Extraction (NDI mean) |
| avg_roi2toroi3 | Tract Extraction (NDI mean) |
| avg_roi1toroi5 | Tract Extraction (NDI mean) |
| avg_roi2toroi6 | Tract Extraction (NDI mean) |
| avg_roi5toroi7 | Tract Extraction (NDI mean) |
| avg_roi6toroi8 | Tract Extraction (NDI mean) |
| avg_roi7topacl | Tract Extraction (NDI mean) |
| avg_roi8topacr | Tract Extraction (NDI mean) |
| avg_roi1toroi3 | Tract Extraction (NDI mean) |
| avg_roi2toroi4 | Tract Extraction (NDI mean) |
| avg_roi3toroi5 | Tract Extraction (NDI mean) |
| avg_roi4toroi6 | Tract Extraction (NDI mean) |
